# Chronic Changes In Oligodendrocyte Sub-Populations After Middle Cerebral Artery Occlusion in Neonatal Mice

**DOI:** 10.1101/2023.01.05.522879

**Authors:** Alexandra P. Frazier, Danae N. Mitchell, Katherine S. Given, Genevieve Hunn, Amelia M. Burch, Christine R. Childs, Myriam Moreno-Garcia, Michael R. Corigilano, Nidia Quillinan, Wendy B. Macklin, Paco S. Herson, Andra L. Dingman

## Abstract

**Background:** Neonatal stroke is common and causes life-long motor and cognitive sequelae. Because neonates with stroke are not diagnosed until days-months after the injury, chronic targets for repair are needed. We evaluated oligodendrocyte maturity and myelination and assessed oligodendrocyte gene expression changes using single cell RNA sequencing (scRNA seq) at chronic timepoints in a mouse model of neonatal arterial ischemic stroke.

**Methods:** Mice underwent sixty minutes of transient right middle cerebral artery occlusion (MCAO) on postnatal day 10 (p10) and received 5-ethynyl-2’-deoxyuridine (EdU) on post-MCAO days 3-7 to label dividing cells. Animals were sacrificed 14 and 28-30 days post-MCAO for immunohistochemistry and electron microscopy. Oligodendrocytes were isolated from striatum 14 days post-MCAO for scRNA seq and differential gene expression analysis.

**Results:** The density of Olig2^+^EdU^+^ cells was significantly increased in ipsilateral striatum 14 days post-MCAO and the majority of oligodendrocytes were immature. Density of Olig2^+^EdU^+^ cells declined significantly between 14 and 28 days post-MCAO without a concurrent increase in mature Olig2^+^EdU^+^ cells. By 28 days post-MCAO there were significantly fewer myelinated axons in ipsilateral striatum. scRNA seq identified a cluster of “disease associated oligodendrocytes (DOLs)” specific to the ischemic striatum, with increased expression of MHC class I genes. Gene ontology analysis suggested decreased enrichment of pathways involved in myelin production in the reactive cluster.

**Conclusions:** Oligodendrocytes proliferate 3-7 days post-MCAO and persist at 14 days, but fail to mature by 28 days. MCAO induces a subset of oligodendrocytes with reactive phenotype, which may be a therapeutic target to promote white matter repair.

## INTRODUCTION

In this study we investigate chronic oligodendrocyte changes after neonatal arterial ischemic stroke. Stroke in the neonatal period is common, with an incidence of 1 in 4,000 live births[2, 3]. Up to 60% of neonates with stroke have long-term motor deficits[4], and at school age 70% have deficits in cognitive ability[5]. In addition, neonatal stroke has significant monetary costs, estimated at over $50,000 per five years[6]. Unlike in adults, neonatal stroke is often not identified within the window of acute thrombolysis or neuroprotection. A little over half of neonates with stroke present at several days of life with focal seizures or encephalopathy, while many infants do not have obvious clinical signs of a perinatal stroke until several months of age[7]. Therefore, a focus on more sub-acute and chronic mechanisms of injury and repair is needed to improve outcome from neonatal stroke. White matter is a potential target for non-acute therapeutic interventions as imaging in children who suffered neonatal stroke suggest that white matter tract injury evolves over several years[8].

White matter injury and repair is mediated by the response of oligodendrocytes (myelin producing cells) to ischemia. While the oligodendrocyte response to ischemia has been studied in models of preterm-brain injury[9], the oligodendrocyte response to focal ischemia, both acutely and chronically, has not been extensively studied in models of full-term neonatal stroke. In preterm brain injury, late oligodendrocyte progenitors are vulnerable after ischemia, but there is also proliferation of a “reactive” oligodendrocyte population[9]. However, myelination and oligodendrocyte maturity changes rapidly between even the late-preterm period and full term, likely contributing to a developmentally variable ischemia response. We previously found that the oligodendrocyte response to stroke depends on age. In juvenile mice (postnatal-day 25) oligodendrocyte progenitor cells (OPCs) increase significantly in the injured striatum 3-7 days post-MCAO, and mature oligodendrocytes are resistant to ischemia in this same time window, with relative preservation of subcortical white matter[10]. In contrast, in adult mice OPCs and mature oligodendrocytes are lost within the injury and the white matter is more severely injured. In a model of focal white matter-specific stroke there is marked proliferation of OPCs after ischemia in juvenile mice but not adult mice, but these newborn cells fail to mature into myelinating cells chronically[11]. These data strongly suggesting that the acute vulnerability and proliferation of oligodendrocytes in response to ischemia is age dependent, and that studying the oligodendrocyte response to stroke in age-relevant models is critical. Therefore, in this study we investigate acute oligodendrocyte injury, proliferation, as well as chronic oligodendrocyte fate in a mouse model of term equivalent (postnatal day 10-11) arterial ischemic stroke.

The main role of oligodendrocytes in the CNS has classically been attributed to myelin sheath formation. However, it is increasingly understood that oligodendrocytes have diverse sub-populations and homeostatic roles beyond myelin formation and maintenance[12, 13]. In addition, recent studies utilizing single cell RNA sequencing (scRNA-seq) have revealed that several pathologic conditions, including multiple sclerosis (MS) and spinal cord injury, are associated with a disease-specific sub-set of oligodendrocytes[1, 14-16]. However, changes in oligodendrocyte populations after ischemic stroke has not been extensively investigated, despite the fact that white matter often comprises a large proportion of the injury area[17]. In the current study, we use scRNA-Seq after MCAO to evaluate gene expression changes in oligodendrocyte sub-populations in response to focal ischemia.

## METHODS

### Animals

All experimental protocols were approved by the Institutional Animal Care and use Committee at the University of Colorado Anschutz Medical Campus and conformed to the National Institutes of Health guidelines for the care and use of animals in research. Male postnatal C57Bl/6 mice were used for all studies (Charles River, USA). Housing conditions were controlled with a 14:10 hour light:dark cycle and all animals received food and water ad libitum.

### Middle Cerebral Artery Occlusion

MCAO or Sham surgery was performed in male postnatal day 10 or 11 (p10-11) mice, as previously described[18]. Timing of surgery is referred to as p10 hereafter for brevity. Littermates were randomly assigned to MCAO or Sham surgery by an independent investigator. Animals underwent anesthesia induction with 3% isoflurane with gas mixture of 20% O2 and 80% compressed air and maintained with 1.5-2% isoflurane. Animal temperature was maintained at 35-37° Celsius during the procedure using a heating pad and monitored with a rectal thermometer. The internal carotid (ICA) and common carotid (CCA) arteries were exposed, and a loose knot was tied at the origin of the ICA. A second ligature was looped just below the ICA origin and retracted laterally to prevent retrograde flow. An arteriotomy was made and rubber-coated 7-0 0.15mm diameter monofilament (Doccol,Sharon, MA, catalog #701512PK5Re) was inserted and advanced 4.5mm to the origin of the middle cerebral artery (MCA). The knot at the ICA origin was then tightened to keep the filament in place. The neck incision was closed and the pups were allowed to recover. Cerebral reperfusion was performed 60 minutes after occlusion when the pups were re-anesthetized, the neck incision was re-opened, the ICA ligatures were removed and the monofilament extracted. After again closing the neck incision pups were returned to their dams. For Sham surgery the ICA was exposed and loosely ligated without occluding for 60 minutes. Animals were either sacrificed on post-MCAO day 3, 14 or 28-30. Animals that were survived to 28-30 days post-MCAO were weaned at p21-25 and singly housed until sacrifice. Litters were left undisturbed for 3 days post-surgery to minimize Dam rejection. Animals were excluded a-priori from the experiment if there was visible intracranial hemorrhage at the time of sacrifice.

### EdU Injections

To determine cell proliferation acutely after WM stroke, animals were injected intraperitoneally with 5-ethynyl-2’-deoxyuridine in sterile saline (EdU, Thermo Fisher Scientific, USA, catalog #A10044), at a dose of 50mg/kg mouse on post-MCAO day 3-7 in single daily injections. EdU concentration was 1mg/mL and injection volume was 5µL/ gram mouse.

### Immunohistochemistry

Animals were sacrificed 3, 14 or 28-30 days post-MCAO. Animals were continuously anesthetized with isoflurane and were transcardially perfused with cold phosphate buffered saline, followed by 4% paraformaldehyde. Brains were then removed, postfixed overnight, and transferred to cryoprotection solution (20% glycerol in 0.1 M Sorenson’s buffer, pH 7.6). Contralateral hemisphere was marked in fixed brains with a shallow razor slice to the cortex. Brains were then frozen, and cut on a sliding microtome in the coronal plane in sections 50 µm in thickness. Serial sections were collected starting at the level of visible lateral ventricles and ending at the last section prior to hippocampus (Allen Adult Coronal MouseBrain atlas plates 44-63, https://mouse.brain-map.org/experiment/thumbnails/100048576?image_type=atlas). Sections were stored in cryostorage solution (30% ethylene glycol, 30% sucrose, and 1% PVP-40 in 0.1 M Sorenson’s buffer). For each combination of markers, two free floating sections 600 µm apart at the level of the striatum (Alan Brain Atlas Coronal Mouse Plates 46-55) were washed in PBS and antigen retrieval was performed with a 10mM tris-EDTA buffer with 0.05% Tween 20 (pH 9) for 10 min at 550 W in a PELCO BioWave Pro tissue processor (Ted Pella, Redding, CA). Sections were then blocked and permeabilized with 5% normal donkey serum (NDS) with 0.3% Triton-X 100. The click-IT EdU kit (ThermoFisher, catalog #C10339) was used to label cells with EdU per manufacturer’s instructions. Sections were then incubated in primary antibodies overnight in 3% NDS, followed by fluorescent secondary antibodies raised in donkey for one hour at room temperature. Primary antibodies used were : rabbit anti-Olig2 (Millipore, USA, Cat# AB9610, RRID:AB_570666), mouse anti-APC/CC1 (Millipore Cat# OP44, RRID:AB_2242783), mouse anti-NeuN (Millipore Cat# MAB377, RRID:AB_2298772), mouse anti-MBP (Abcam, Eugene, OR Cat# ab40390m RRID: AB_1141521), rabbit anti-SMI34 (BioLegend, San Diego, CA, Cat# 835504, RRID: AB_2728546), rat anti-MHCI (Abcam Cat# ab15681), rabbit anti-Gst-pi (Enzo Life Sciences, Farmingdale, NY, Cat#ADI-MSA-102-E).

### Cresyl Violet Staining and Injury Measurement

To evaluate injury volume 3-days post-injury, 6 sections at 300 µm intervals were stained with Cresyl Violet. Briefly, sections were mounted on charged slides and dried. Sections were rehydrated through a series of decreasing Ethanol solutions before incubation for 15 minutes in 0.1% Cresyl Violet solution. Sections were then dehydrated with sequentially increasing Ethanol solutions followed by Xylene. Slides were cover-slipped with DPX mounting media (Sigma Aldrich, Cat #44581). Injury volume was measured using Stereologer Software (www.stereologyresourcecenter.com, RRID:SCR_003137).

### Fluoro-JadeB/NeuN Co-Staining

In order to assess cell death, co-staining with the cell death stain Fluoro-JadeB (FJB)[19] and fluorescent IHC with antibodies for NeuN was performed. Two sequential free-floating sections 600 µm apart were stained with mouse anti-NeuN as above. Sections were then mounted on charged slides and dried. They were they rehydrated through a series of decreasing Ethanol solutions, followed by incubation om 0.06% Potassium Permanganate for 15 minutes. Sections were then incubated in 0.001% Fluoro JadeB (Sigma Aldrich Cat# AG310) for 20 minutes. Sections were then dehydrated through increasing Ethanol solutions to Xylene, and cover-slipped with DPX media.

### Cell Counting

Stained sections were mounted on slides and imaged on an Olympus FV1000 laser scanning confocal microscope using Fluoview software (Olympus Fluoview FV10-ASW, RRID:SCR_014215). For oligodendrocyte counts Images from ipsilateral and contralateral striatum were obtained using a 20x objective with 2x digital zoom, resulting in images with dimensions 317.44 x 317.44 µm^2^. Cells were counted in two non-overlapping images per hemisphere in lateral striatum in two sections per animal, and counts were averaged over the 4 images per hemisphere for each animal. FJB/NeuN counts (Figure 1), were obtained from two non-overlapping images in striatum and in the adjacent cortex using a 20x objective and 1x digital zoom, resulting in images 635.9 x 635.9 µm^2^. For 3D cell reconstruction (figure 8) Imaris Software v 9.8.2 was used.

**Figure 1:**
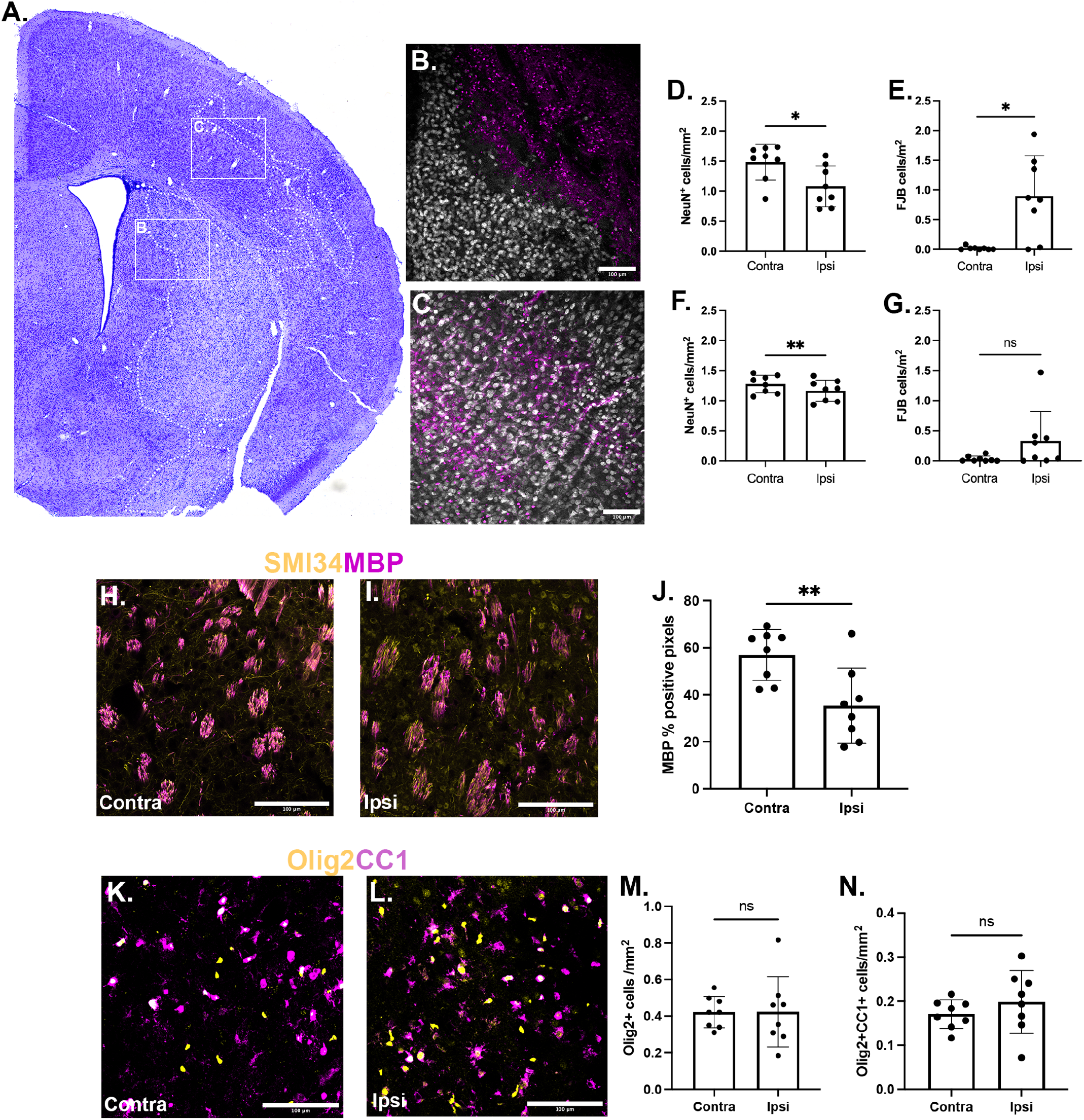
60 min MCAO causes striatal cell death, neuronal and myelin loss 3 days post-MCAO, but not axonal or mature oligodendrocyte loss. P10 mice underwent 60min MCAO followed by 3 days of reperfusion. **A:** Representative coronal section from ipsilateral hemisphere stained with Cresyl Violet. Injury was identified by groups of pyknotic nuclei (white dashed line). **B-C:** Neuron and dying/dead cell density was assessed in striatum (B) and cortex (C) with staining with Flouro-Jade B (magenta) and antibodies to NeuN (gray). **D-E:** Quantification of NeuN^+^(D) and FJB^+^ (E) cell density in striatum. **F-G:** quantification of NeuN^+^(F) and FJB^+^(G) cells in cortex. **H-I:** Immunofluorescence to SMI34(magenta) and MBP (yellow) was used to label axons and myelin respectively in contralateral (H) and ipsilateral (I) striatum. J: Quantification of % positive MBP pixels within pencil fibers (defined by SMI34^+^ axon groups). **K-L:** Brain sections were incubated with antibodies for Olig2 (K&L, yellow) to identify oligodendrocytes and CC1 (K&L, magenta) to label mature oligodendrocytes. **M-N:** Quantification of Olig2^+^ cells (M) and Olig2^+^CC1^+^ cells(N) in ipsilateral and contralateral striata. *p<0.05, **p<0.01, paired t-test.

### Fluorescent Myelin Analysis

To assess acute changes in myelin 3-days post-injury, sections were stained with antibodies to axons (SMI34_ and myelin (MBP) and imaged using a confocal microscope (20x objective with 2x digital zoom, dimensions 317.44 x 317.44 µm^2^). To assess myelin content specifically in pencil fibers, regions of interest (ROIs) were first drawn around radial fiber bundles based on SMI34 staining after applying a standard image threshold across all images. The SMI34-based ROIs were then applied to thresholded MBP images and % MBP positive pixels within ROIs were measured.

### Electron Microscopy and Pencil Fiber Evaluation

Animals underwent MCAO at p10-11 and were anesthetized at 14 or 28-30 days post-MCAO and perfused with fixative containing 2% paraformaldehyde, 2.5% glutaraldehyde, 2mM calcium chloride and 0.1M cacodylate buffer, pH 7.4. The brain was stored in postfix until embedding. Striatal tissue just medial to the corpus callosum was isolated from 300-µM coronal slices. Tissue was postfixed in 1% osmium tetroxide, dehydrated in graded acetone, and resin embedded in Embed 812 (Electron Microscopy Services, Hatfield, PA, Cat# 14120). Striatal pieces were oriented such that sections could be cut in a coronal plane to visualize myelinated axon fibers. Ultrathin sections (80nm) were mounted on copper grids, stained with uranyl acetate and lead citrate, and viewed at 80 kV on a Tecnai G2 transmission electron microscope (FEI Company).

G-ratios of myelinated axons from striatal radial fibers (pencil fibers) were calculated as the ratio of the diameter of the axon to the diameter of the myelinated fiber. Diameters were derived by measuring the respective perimeters using FIJI software. A minimum of 100 axons from at least 6 radial fiber bundles were counted from ipsilateral and contralateral lateral striatum from two sections per animal. The density of myelinated axons per field of view was also quantified. Myelinated axons with pathologic features were counted using Fiji Software, and the ratio of pathologic to all myelinated axons was calculated. Pathologic features were defined as vesicles in the inner or outer myelin tongue or myelin protrusions.

### Striatal Cell Dissociation

Oligodendrocytes were isolated for single cell RNA sequencing from 3 animals at 14 days post-MCAO. Mice were transcardially perfused with ice-cold (2-5°C) oxygenated (95% O_2_/5% CO_2_) artificial cerebral spinal fluid (aCSF) for two minutes prior to decapitation. The brains were then extracted and placed in the same aCSF. The aCSF was composed of the following (in mmol/L): 126 NaCl, 2.5 KCl, 25 NaHCO_3_, 1.3 NaH_2_PO_4_, 2.5 CaCl_2_, 1.2 MgCl_2_ and 12 glucose. Whole brains were trimmed of the cerebellum and hemi-sected in the sagittal plane to separate ipsilateral and contralateral hemispheres. Coronal slices from each hemisphere (300μm thick) were cut with a Vibratome 1200 (Leica). Lateral striatum was dissected from each hemi-section and placed in HABG media (Hibernate A Media, BrainBits, Springfield, IL, HA500; 2% B27 Supplement, ThermoFisher, Cat #17504044; 0.26% Glutamax, ThermoFisher, Cat# 35050061). For each sample (ipsilateral and contralateral), striata from 3 animals was pooled in the same sample for biologic variability. Tissue was washed by centrifuging in a swinging-bucket table-top centrifuge (Beckman Coulter Allegra X-14R) at 400xg. The tissue was resuspended in 3 mL of 3.5% Collagenase/Dispase (Sigma, USA, 11097113001, 100mg/mL stock) in Hibernate A minus Calcium Media (BrainBits, HACA100) with 0.125% Glutamax. Samples were then dissociated using the GentleMacs Octo Dissociator (Miltenyi Biotech, Githersburg, MD, 130-195-937) on program 37_NDK_01. After dissociation enzymes were quenched with HABG up to 15 mL. Tubes were spun for 5 minutes at 400xg. The tissue was resuspended in 2 mL HABG media, and 50 uL of Deoxyribonuclease I from bovine pancrease (DNASE) was added. Each sample was triturated about 10 times with a flame polished glass pipette. Dissociated tissue was then passed through a 100µm cell strainer (Falcon, 352360), followed by a 40µm cell strainer (Falcon, 352340). Each strainer was washed with 20mL of HABG media. Samples were then spun for 5 minutes at 400xg and resuspended in 4mL of 30% Percol (Sigma, P1644) in Calcium- and Magnesium-free Hank’s Balanced Salt Solution (HBSS, ThermoFisher, 14170112). One milliliter of HBSS free of calcium and magnesium was then carefully layered on top of the Percol, and layers were allowed to separate for 5 minutes on ice. Samples were then centrifuged for 15 minutes at 700xg with the brake turned off. The top layer of myelin and cellular debris was discarded, and the remaining layer was washed with 15mL of HABG media. Following centrifugation at 400xg for five minutes, the cell pellet was used for fluorescent activated cell sorting (FACS).

### Flow Cytometry and Fluorescent Activated Cell Sorting

Preliminary experiments determined the antibodies and titrations to capture the full breadth of oligodendrocyte lineage cells. The following fluorescently conjugated antibodies were tested and cell analysis was done on a Bio-Rad ZE5/Yeti cell analyzer: APC-A2B5 1:400 (Miltenyi Cat # 130-098-039), PE-O4 1:250(Miltenyi Cat #130-117-507), and FITC-O1 1:400 (R&D Biosciences, Minneapolis, MN, Cat# FAB1327G). BV421-CD45 1:500 (BD Biosciences, Franklin Lakes, NJ, Cat# 5638-90) antibodies were used to label immune cells. Flow buffer consisted of 2% Fetal Bovine Serum; 1X DPBS without Ca or Mg (Sigma) and was the diluent for all blocking and immunofluorescent antibody staining. Cells were blocked with 1.2% Fc block (BD Biosciences, Cat# 553141) for 10 minutes at 4°C, and then incubated in primary antibodies for 30 minutes at 4°C. Dapi(Sigma Cat # D9542) was added to cells just before analysis to label dead and dying cells. We found near complete overlap of O1 and O4 antibodies (supplemental figure 1A). Therefore, we used A2B4 and O4 antibodies for final cell sorting to collect both oligodendrocyte precursors and oligodendrocytes for RNA-seq. Cell sorting was done with a SonyMA900 cell sorter. For final cell sorting unstained and fluorescence minus one (FMO) controls were included. Ultracomp eBeads (ThermoFisher, 01-2222-41) were used for single antibody controls. Scatter was used to gate for non-debris singlet cells (supplemental figure 1B&C). Cells that labelled with Brilliant Violet 421 (BV421)/Dapi (ie CD45^+^ or dead/dying cells) were excluded (supplemental figure 1D), and remaining cells that were A2B5+ and/or O4+ (ie anything positive, supplemental figure 1E) were sorted for RNA-seq.

### Single Cell RNA Sequencing and Data Analysis

Single Cells were captured using the 10x Genomics Single Cell 3’ Reagent Kit v3 on an Illumina NovaSEQ 6000 instrument. Illumina basecall files were converted to FASTQ files and were processed using CellRanger v3.1.0 and aligned to the mouse reference genome mm10 (v3.0.0) as previously described [20]. The mean number of reads per cell was 252,000. QC and downstream processing was done using Seurat version 4.0.1 run on R Studio version 1.4.1106 using R version 4.1.1 [21]. Quality Control (QC) filtering removed cells that had less than 400 or greater than 6000 features and greater than 15% expression of mitochondrial genes. After QC filtering our data consisted of 1,833 cells from the contralateral hemisphere and 2,317 from the ipsilateral hemisphere. Initial cell clustering was performed using the SCT transformation modeling framework built into Seurat and previously published[22]. Cell types were determined by comparing gene expression to the Tabula Muris mouse transcriptome database[23], revealing 6 CNS cell types (supplemental figure 2A). Non-oligodendrocyte lineage cells were excluded for the remainder of analysis. Oligodendrocyte lineage cells were re-clustered, resulting in 9 subsequent cell clusters (supplemental figure 2B-C). We screened all cell clusters for expression of the ubiquitous oligodendrocyte genes, Olig2 and Sox10 (supplemental figure 2D). Two cell clusters were excluded due to low percentage of cells expressing these genes (clusters 5&6, supplemental figure 2D), resulting in 7 final cell clusters for downstream analysis (supplemental figure 3A). Pseudo time analysis was done using Monocle3 with a Seurat wrapper[24], using the oligodendrocyte progenitor predominant gene Pdgfra as the anchor. A phylogenetic tree of all cell clusters was built, and differential gene expression followed by gene ontology analysis was performed at each node of the phylogenetic tree using G:Profiler[25]. Differential gene expression (DE) analysis was performed using Seurat.

### RNA-Scope Fluorescent In Situ Hybridization

Animals underwent 60-minute MCAO followed by reperfusion and were sacrificed with PFA perfusion-fixation 14 days later. Brains were stored in cryoprotection solution prior to freezing and cutting to 50µm thick sections and stored in cryostorage buffer made with RNAase free water. To detect B2M, Neat1, and SOX10 mRNA from oligodendrocyte populations in fresh frozen striatal sections from neonatal MCAO mice, fluorescent *in situ* hybridization (FISH) was performed using RNAscope® Multiplex Fluorescent Reagent Kit (cat# 320293, Advanced Cell Diagnostics) according to the protocol provided by the manufacturer. Briefly, microtome-cut 50µM frozen sections were mounted on charged slides, air dried, and fixed in 4% paraformaldehyde buffer. Following fixation, sections underwent a series of ethanol dehydration steps (50, 70, 80, 100%) and were heated in manufacturer-provided antigen retrieval buffer. Sections were then subjected to protease (kit-provided) digestion for 30 minutes at 40°C. For hybridization, sections were exposed to probes (C1 B2M cat# 415191, C2 Neat1 cat# 538761-C2, C3 SOX10 cat#435931-C2; Advanced Cell Diagnostics) and incubated at 40°C in a hybridization oven for 2 hours. Following washes, sections underwent signal amplification steps and fluorophore (1:750 dilutions of Opal™ Dyes: 520 cat# FP1487001KT, 570 cat# 1488001KT, 690 cat# 1497001KT; Akoya Biosciences) treatments according to manufacturer’s instructions. Sections were counterstained with DAPI (kit-provided), stored at 4°C and imaged within two weeks. Images were acquired from lateral striatum in the ipsilateral and contralateral hemispheres from two neonatal MCAO mice using a 20x objective with 2x digital zoom, resulting in images with dimensions 317.44 x 317.44 µm^2^.

### Statistics and Rigor

All experiments were performed in accordance with the Arrive 2.0 guidelines[26]. For 3-day IHC analysis the contralateral hemisphere served as control. For chronic time-points IHC analyses were done in pups that underwent MCAO or sham surgery. MCAO and sham surgeries were performed in litter-mates, randomized by an independent investigator. Because there was no difference in numbers of OPCs or mature oligodendrocytes in contralateral vs sham hemispheres, for EM and scRNAseq contralateral hemisphere served as control. Animal identification in regards to type of surgery was blinded for image acquisition and hemisphere was blinded for cell counting. Graphs and statistics were generated on Graphpad Prism 9 for IHC analyses, and the lmer package in R for electron microscopy analyses. Normality criteria was determined per group and data set using the Shapiro-Wilks’ test. Analyses involving two groups were analyzed using paired t-test(parametric) or Wilcoxon test (non-parametric). When more two or more conditions were compared 2-way ANOVA or mixed effects model was used. Data is reported in the text as mean±SD unless otherwise stated. Complete statistical analyses of all comparisons are available in supplementary table 1.

For EM g-ratio, axon density and abnormal myelin morphology analyses, because many pencil fibers and axons were analyzed over a small number of animals (n=4 per time point), we used a general linear mixed model to assess the difference in myelinated axon density or g-ratio as a function of hemisphere (contra vs ipsi) and timepoint (14d vs 28d post-MCAO), with mouse ID as the random effect. Because response variables (axon density, g-ratio, and ratio abnormal axons) are values from 0-1, a beta distribution with logistic link function was used. Myelinated axon density and g-ratio respectively were modeled as a function of hemisphere, timepoint, and mouse ID (random effect) with the following equations:

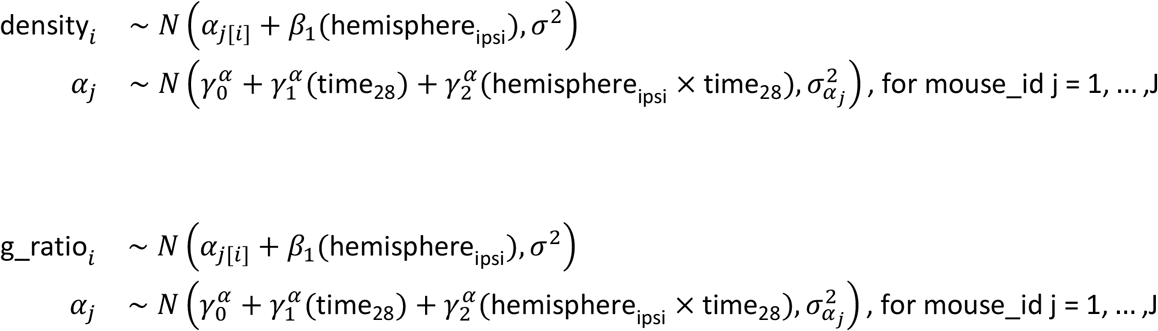

Ratio of number of axons with abnormal myelin structure to total number of myelinated axons was modeled as a function of hemisphere and mouse ID (random effect) at 28 days post-injury with the following equation:

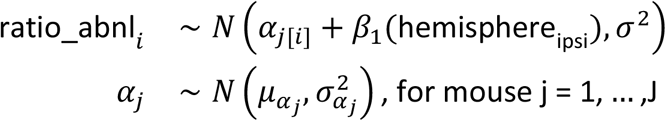

General linear mixed modeling was performed using the lmer4[27] package in R version 4.1.1. Post-hoc analysis was done with the emmeans[28] package in R, using Tukey’s adjustment for multiple comparisons. Complete statistics of all comparisons are available in supplementary table 2. All statistical analyses for scRNAseq data was done per the default tests built into the Seurat package for R[22].

## RESULTS

### Survival

For long-term survival studies (14- and 28-days post-surgery), a total of 37 mice were randomized to MCAO and 26 to Sham surgery. Mortality was 43% in the MCAO group. There was no mortality in the Sham group. Litters in which all pups died (including Sham and non-operated pups) were excluded from mortality calculation as this most often occurred with inexperienced dams and is assumed to be due to litter rejection. A total of 5 pups in the MCAO group and 2 in the Sham group were excluded due to evidence of intracranial hemorrhage at the time of sacrifice.

### 60-minute MCAO causes acute neuronal loss in p10-11 mice

To characterize the cellular response to injury following neonatal ischemic stroke, postnatal day 10 mice were subjected to 60 min MCAO and brains were analyzed by histology and immunohistochemistry 3 days after reperfusion. Median injury volume measured in Cresyl Violet (CV) stained 14sections was 10.36% of ipsilateral hemisphere volume (IQR 2.67-18.48%). One out of eight animals analyzed had an injury volume of 0%. A representative CV-stained coronal section with injury outlined is shown in figure 1A. Injury was defined as clusters of pyknotic nuclei. Increased cell death, assessed with Fluoro-Jade B (FJB) positive cells, and neuronal loss, assessed by NeuN^+^ cell density, was observed in the ipsilateral striatum (figure 1B) and cortex (figure 1C), compared to the non-ischemic contralateral. Cell death and neuronal loss were greater and more consistent in ipsilateral striatum (FJB^+^ Contra: 0.0184 ± 0.028 vs FJB+ Ipsi: 0.895 ± 0.681 cells/mm^2^ p=0.02, NeuN^+^ Contra:1.484 ± 0.300 vs NeuN+ Ipsi:1.078 ± 0.340 cells/mm^2^ p=0.04, figure 1 D-E). The magnitude of cell death and neuron loss was less in adjacent cortex (FJB^+^ Contra: 0.031 ±0.045 vs FJB^+^ Ipsi: 0.331 ± 0.491 cells/mm^2^ p=0.05, NeuN^+^ Contra: 1.277 ±?0.147 vs NeuN^+^ Ipsi: 1.166 ± 0.156 cells/mm^2^ p=0.003, figure 1F-G) compared to striatum.

### Acute Changes in Striatal White Matter and Oligodendrocytes

We next determined whether myelin and mature, myelinating oligodendrocytes were lost in the same region of acute neuronal loss at 3 days post-MCAO. Myelin within striatal pencil fibers at this timepoint were assessed with immunofluorescence labeling using antibodies to myelin basic protein (MBP) and the neurofilament protein SMI-34. Figure 1H-I demonstrates that striatal radial white matter tracts (aka pencil fibers) are grossly preserved in the injured area. However, within pencil fibers (defined by groups of SMI^+^ axons) the density of MBP^+^ myelin was significantly decreased (contra: 56.90 ± 10.77 vs ipsi: 35.36 ± 16.03 % MBP^+^ pixels, p= 0.003, figure 1J). Despite the acute loss of myelin, oligodendrocytes in neonatal striatum appear to be highly resistant to ischemic injury. We assessed the density of all oligodendrocytes (Olig2^+^) and mature oligodendrocytes (Olig2^+^CC1^+^) in lateral striatum 3-days post-injury. The CC1 antibody binds Quakin7, a protein upregulated in myelinating oligodendrocytes[29]. Because CC1 can also be expressed in a sub-set of astrocytes[30], only CC1+ cells that also expressed Olig2 were included in analysis. There was not a significant decrease in the density of total oligodendrocytes (Contra Olig2^+^: 0.422 ± 0.086 vs Ipsi: 0.424 ± 0.192 cells/mm^2^, p = 0.98, figure 1M) or in mature oligodendrocytes (Contra Olig2^+^CC1^+^: 0.171 ± 0.033 vs Ipsi: 0.199 ± 0.071 cells/mm^2^, p=0.35, figure 1N) in ipsilateral striatum compared to contralateral.

### Oligodendrocytes proliferate sub-acutely after p10 MCAO and persist 14 days post-MCAO

Previous studies have shown that oligodendrocytes proliferate 3-7 days after ischemia in young animals[10, 11]. Therefore, the thymidine analog, EdU, was injected into animals on days 3-7 post-MCAO to label dividing cells (figure 2A). Striatum ipsilateral and contralateral to MCAO was then assessed 14 and 28 days post-MCAO for EdU positive oligodendrocytes (Olig2^+^). At 14 days post-MCAO there were many more Olig2^+^ cells in ipsilateral striatum compared to contralateral (figure 2B&E). In addition, there were many more EdU^+^ cells in ipsilateral striatum (figure 3C&F), and the majority of oligodendrocytes in ipsilateral striatum were EdU^+^ (figure 2G), indicating their birth 3-7 days after ischemia. Overall, there was a significantly higher density of EdU^+^ oligodendrocytes (figure 2J) in ipsilateral striatum compared to contralateral striatum (contra: 0.03 ± 0.02 vs ipsi: 0.38 ± 0.37 Olig2^+^EdU^+^ cells/mm^2^, p=0.001) and compared to sham (sham ipsi: 0.06 ± 0.03 Olig2^+^EdU^+^ cells/mm^2^, p=0.0002).

**Figure 2:**
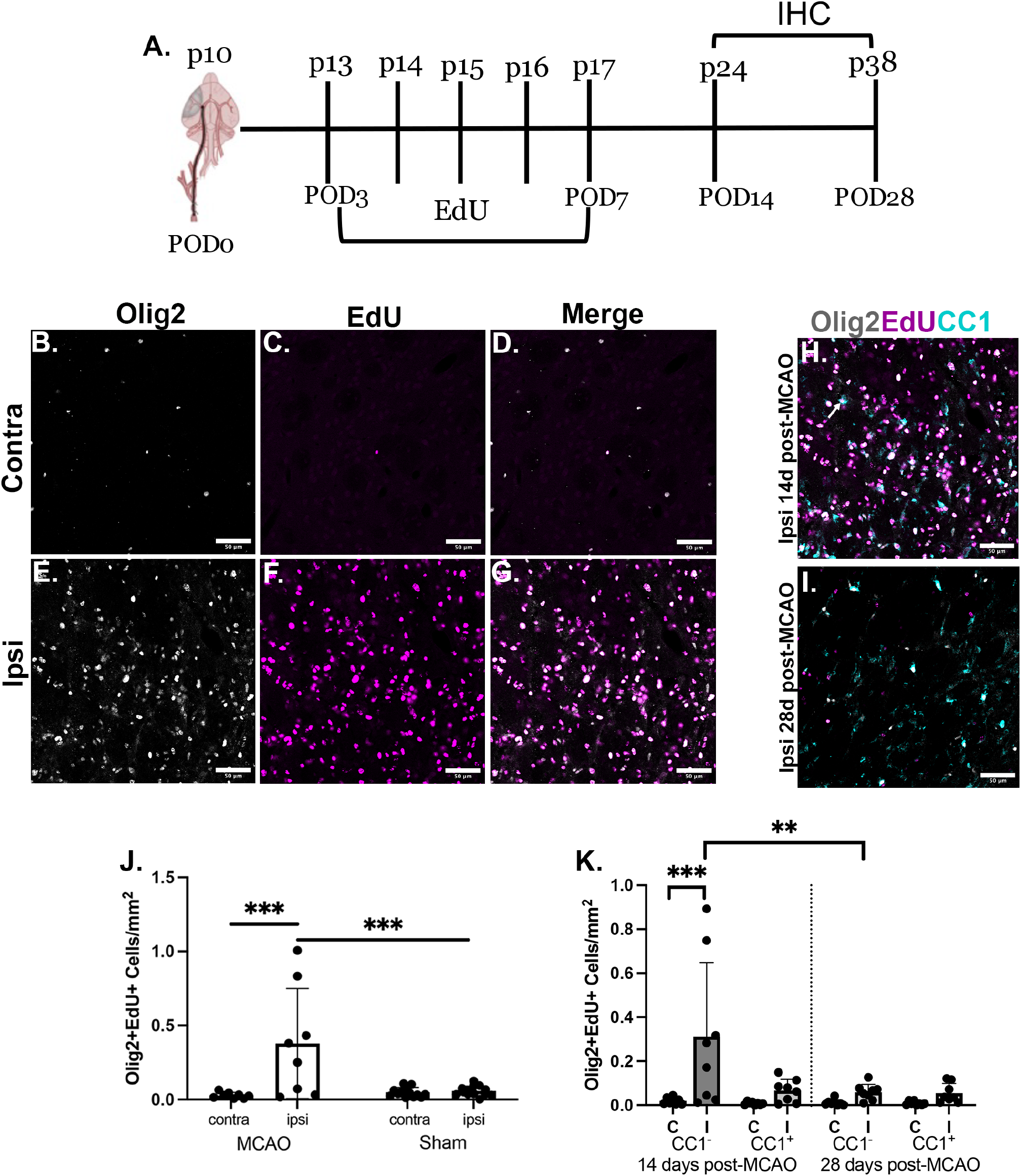
OPC’s proliferate on POD 3-7 and persist as immature OLs on POD 14, but not as mature OLs on POD 28. **A:** Experimental timeline. **B-G:** Immunofluorescence for the oligodendrocyte marker Olig2, and EdU in contralateral (B-D) and ipsilateral (E-G) lateral striatum at 14 days post-MCAO. **H-I:** Immunofluorescence for Olig2, EdU, and the mature OL marker CC1 in ipsilateral striatum at different timepoints. At 14 days post-MCAO (H) few Olig2^+^Edu^+^ cells co-expressed the mature oligodendrocyte marker, CC1 (arrow). By 28 days post-MCAO (I) very few Edu^+^Olig2^+^ cells remain, and there are few triple positive cells. **J:** Quantification of Olig2^+^EdU^+^ cells at 14 days post-MCAO. ***p<0.001, two-way ANOVA. K: Quantification of immature (CC1^-^) and mature (CC1^+^) EdU^+^ oligodendrocytes at 14 and 28 days post-MCAO *p<0.05, **p<0.01, mixed effects model).

**Figure 3:**
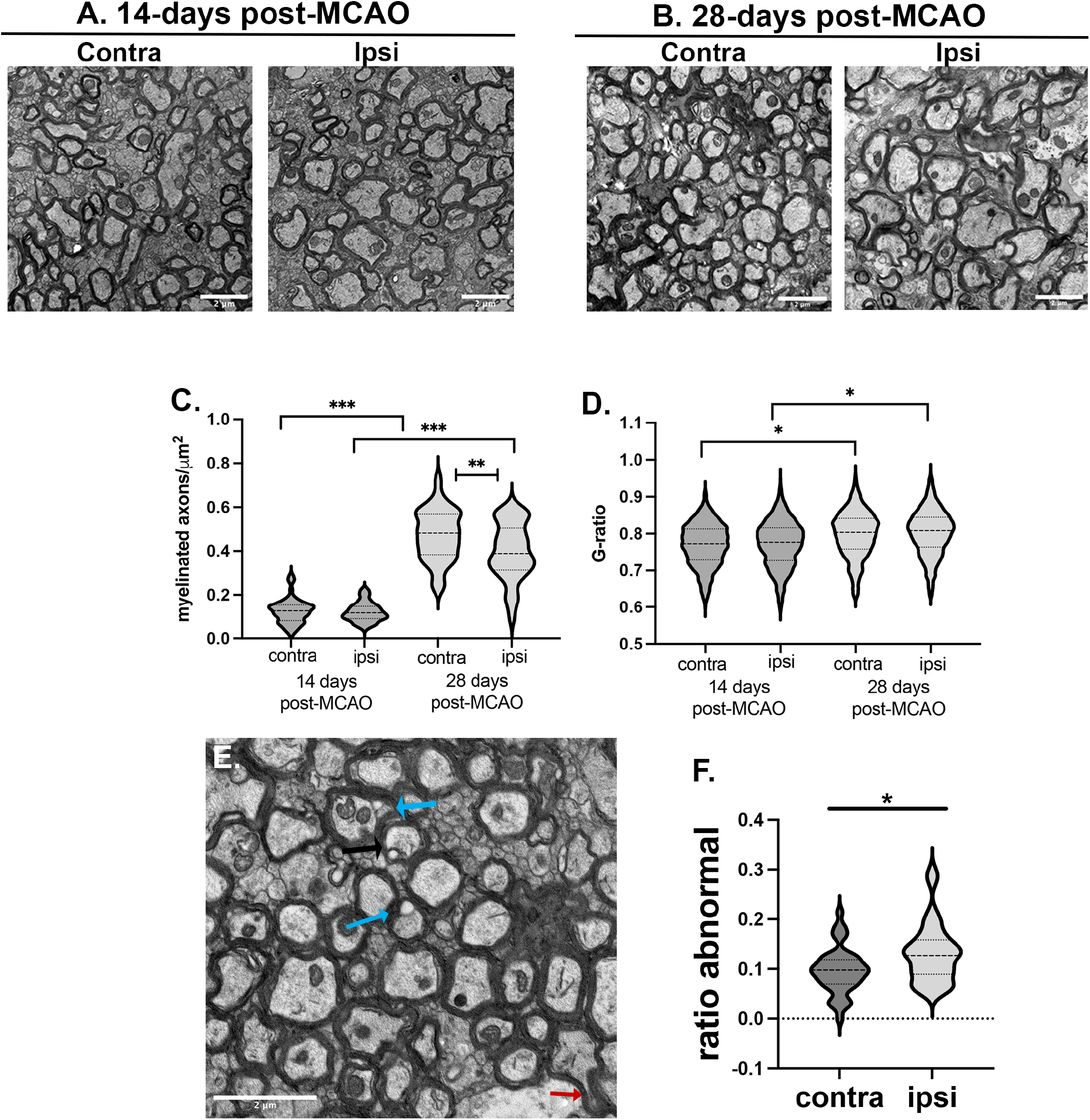
Change in myelination between 14 and 28 days post-MCAO. **A-B:** Electron microscopy(EM) images from lateral striatum 14 (A) and 28 days (B) post-MCAO. **C-D:** Quantification of myelinated axon density (C) and g-ratio (D) 14 & 28 days post-MCAO. **E:** representative image from ipsilateral striatum 28-days post-injury shows inner tongue vesicles (blue arrows) and myelin protrusions (red arrow). **F:** Quantification of axons with abnormal myelin features, expressed as a ratio to all myelinated axons. * p<0.05, ** p<0.01, ***p<0.001, pairwise comparisons of estimated marginal means using a generalized linear mixed model with Tukey’s correction for multiple comparisons.

### Newborn oligodendrocytes remain immature at 14 days post-MCAO but do not mature by 28-days post-MCAO

In order to determine the maturity of newborn oligodendrocytes at 14 and 28 days post-MCAO, we co-stained tissue with the mature oligodendrocyte marker, CC1, to distinguish between immature (CC1^-^) and mature (CC1^+^) cells. Figure 2H-I shows a comparison of immature vs mature oligodendrocyte density in the ipsilateral striatum between 14 and 28 days post-MCAO. At 14 days post-MCAO there is a dramatic increase in immature (Olig2^+^EdU^+^CC1^-^) oligodendrocytes in ipsilateral striatum compared to contralateral (contra: 0.020 ± 0.015 vs ipsi: 0.312 ± 0.337 Olig2^+^EdU^+^CC1^-^ cells/mm^2^, p=0.0002), but not in mature EdU^+^ oligodendrocytes (contra: 0.008 ± 0.007 vs ipsi: 0.066 ± 0.051 Olig2^+^EdU^+^CC1^+^ cells/mm^2^ p=0.995, figure 2K). Between 14 and 28 days post-MCAO there was a significant decrease in the density of immature oligodendrocytes in ipsilateral striatum (14 days: 0.312 ± 0.337 vs 28 days: 0.059 ± 0.036 Olig2^+^EdU^+^CC1^-^ cells/mm^2^, p= 0.0014). If the decrease in immature oligodendrocytes between these timepoints were due to their maturation into mature cells, we would expect the number of CC1^+^ cells to *increase* in ipsilateral striatum between 14 and 28 days post-MCAO. However, mature (CC1^+^) oligodendrocyte density did not change between 14 and 28 days post-MCAO (14 days: 0.066 ± 0.051 vs 28 days: 0.054 ± 0.044 Olig2^+^EdU^+^CC1^+^ cells/mm^2^, p>0.9999). Neither the density of CC1^-^ or CC1^+^ EdU^+^ oligodendrocytes were significantly different between contralateral and ipsilateral hemispheres at 28 days post-MCAO (p=0.9990 and 0.9994, respectively). There was no difference in CC1- or CC1+ Olig2EdU+ cells between contralateral and sham groups at any timepoint (supplemental figures 3A-D).

### Myelin changes between 14 and 28 days post-MCAO

Ischemia has been shown to affect the structure of myelin[31], although we have previously shown that myelin structure is relatively preserved in juvenile mice compared to adult[10]. We assessed the lateral striatum pencil fibers using electron microscopy, and measured the density of myelinated axons *within* pencil fibers, as well as the g-ratio of individual axons (figure 3 C&D). In 4 experimental mice, a total of 4,738 axons in 181 pencil fibers (14-day contra: 838 axons in 60 pencil fibers, 14-day ipsi: 719 axons in 54 pencil fibers, 28-day contra: 1693 axons in 33 pencil fibers, 28-day ipsi:1488 axons in 34 pencil fibers) were analyzed. Because there was no significant difference in oligodendrocyte proliferation or maturation between MCAO contralateral and Sham groups, EM analyses was only performed after MCAO and comparisons between contralateral and ipsilateral hemispheres were made. At 14 days post-MCAO there was not a significant difference in the density of myelinated axons within pencil fibers between contralateral and ipsilateral striatum (contra: 0.113±0.016 vs ipsi 0.116±0.015 myelinated axons/µm^2^, mean±SEM, p=0.9983, figure 3C). However, there was a significant increase in the density of myelinated axons within pencil fibers between 14- and 28-days post-MCAO in both contralateral (14 day: 0.113±0.015 vs 28 day: 0.470±0.017 myelinated axons/µm^2^, p<0.0001), and ipsilateral (14 day: 0.116±0.015 vs 28 day: 0.399±0.017 myelinated axons/µm^2^). Although there was an increase in myelinated axons in both hemispheres over the course of the study, at 28-days post-MCAO there were significantly fewer myelinated axons in ipsilateral striatum compared to contralateral (contra 28 day: 0.470±0.017 vs ipsi: 0.399±0.017 myelinated axons/µm^2^, p=0.0025). The g-ratio is a ratio of axon diameter to the diameter of the myelinated fiber (axon+myelin sheath). Therefore, g-ratio is inversely proportional to myelin thickness. There was not a difference in g-ratio between contra and ipsi hemispheres at 14 days post-MCAO (contra: 0.771±0.005 vs ipsi: 0.772±0.005, p=0.9976, Figure 4C) or at 28 days post-MCAO (0.798±0.005 vs 0.803±0.005, p=0.1036). There was a small, but significant increase in g-ratio over the course of the study in contralateral(14day: 0.771±0.005 vs 28 day: 0.798±0.005, p=0.0288) and ipsilateral(14 day:0.772±0.005 vs 28 day:0.803±0.005, p=0.0142). We also assessed morphologic features of myelin pathology at 28-days post-injury, specifically the presence of inner and outer tongue vesicles and myelin protrusions (figure 3E), both of which have been described in experimental models of abnormal myelinaiton [32, 33]. Although there was no difference in g-ratio in myelinated pencil-fiber axons at 28-days post-injury, there was a significantly greater ratio of axons with abnormal pathologic features within myelin in ipsilateral striatum (Contra ratio abnormal axons: 0.097±0.013 vs Ipsi: 0.129±0.015, p=0.02, figure 3F).

**Figure 4:**
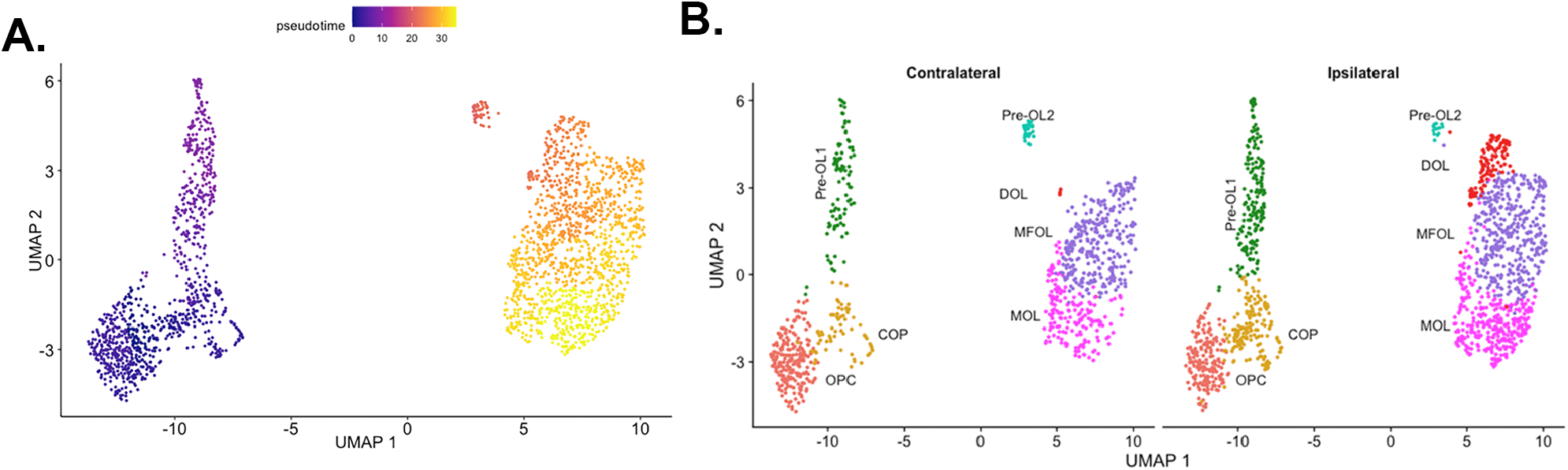
Oligodendrocyte sub-types isolated from mouse striatum. **A:** Pseudo time analysis of oligodendrocytes with Pdgfra as the anchor for oligodendrocyte pre-cursor cells. **B:** Final oligodendrocyte cell clusters separated by hemisphere.

### scRNA-Seq of Oligodendrocytes 14 days post-MCAO

In order to investigate the transcriptional basis for oligodendrocyte maturational failure, we performed single cell RNAseq on oligodendrocytes from striatum ipsilateral and contralateral to MCAO. We chose 14-days post-MCAO because of the significant increase in EdU^+^Olig2^+^CC1^-^ cells at this timepoint. An initial 2,066 cells from pooled contralateral and 2,775 cells from pooled ipsilateral were isolated from lateral striatum and sequenced. After quality control and filtering out non-oligodendrocytes (supplemental figure 2), there were a total of 892 cells from contralateral striatum and 1,386 cells from ipsilateral striatum. Supplemental figure 4A shows a dimensional reduction 2D scatterplot of the 7 final cell clusters, as well as a phylogenetic tree representing the relationship of each cell cluster based on differential gene expression in each cluster (supplemental figure 4B). Pseudo time analysis with Pdgfra as the anchor for least mature cells showed a maturational progression through the cluster plot (figure 4A). Based on the pseudo time plot as well as differential gene expression and gene ontology analysis between each branch in the phylogenetic tree dendrogram (supplemental figure 4C) and relative expression of known maturational markers (supplemental figure 4D), we assigned cell clusters the following identities: oligodendrocyte precursor cell (OPC), committed precursor cells (COP), Pre-Oligodendrocyte 1 and 2 (pre-OL1/2), myelin forming oligodendrocyte (MFOL), mature oligodendrocyte (MOL) and disease associated oligodendrocyte (DOL) (figure 4B). The oligodendrocyte precursor genes *Pdgfra* and *Cspg4* were highly expressed in clusters one and two, but *Bcas1* expression was more consistently expressed in cluster two (supplemental figure 3D), suggesting a commitment to oligodendrocyte maturation [34]. Therefore, we designated clusters one and two OPC and COP, respectively. Cells in clusters three and four had increased expression of *Bcas1 and Tns3*, genes upregulated in newly formed, pre-myelinating oligodendrocytes (supplemental figure 3D) [34, 35]. Therefore, we designated these clusters pre-OL1 and 2 respectively. Both clusters 6 and 7 expressed the more mature oligodendrocyte gene *Plp1*, but *Pmp22*, which is known to be expressed earlier than *Plp1 [36]*, had increased expression in cluster6 (supplemental figure 3D). Furthermore, cluster 6 had increased enrichment in gene ontogeny pathways involved in active myelination, but these pathways were not prominent in cluster 7 (supplemental figure 3C). Therefore, we termed clusters 6 and 7 MFOL and MOL, respectively. We termed cluster 5 “disease associated OL (DOL)” because this cluster was almost exclusively present in the striatum ipsilateral to MCAO (figure 4B). Table 1 shows the proportion of oligodendrocytes in each cluster from contralateral and ipsilateral striata. There was a significant decrease in the proportion of oligodendrocytes in the OPC cluster (25.2% of all oligos in contralateral vs 13.1% in ipsilateral, residual -7.35, p<0.0001, table 1), and in the pre-OL2 cluster (3.5% in contralateral vs 1.0% in ipsilateral, residual - 4.13, p<0.0001). In contrast, there was a significantly *increased* number of cells in the COP cluster (7.7% of all oligos in contralateral vs 11.8% in ipsilateral, residual 3.10, p=0.0271) and in the DOL cluster (0.4% in contralateral vs 8.8% in ipsilateral, residual 8.51, p<0.0001).

**Table 1:**
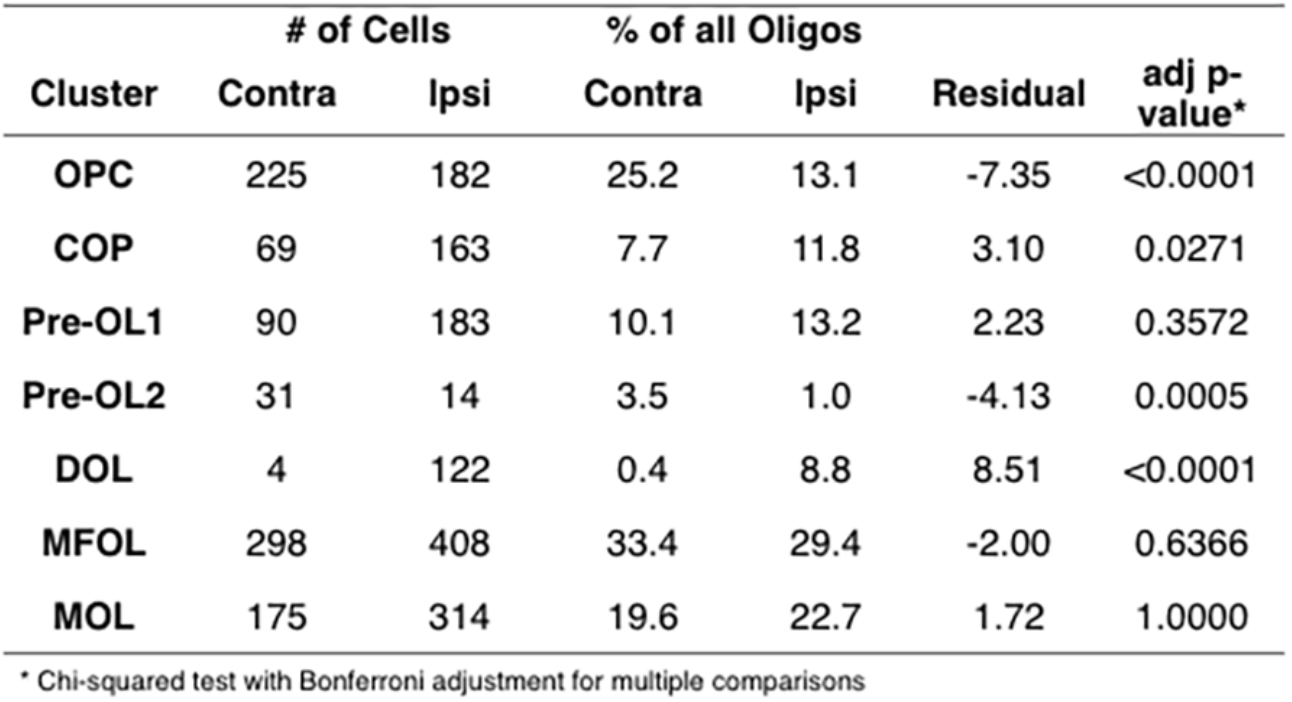
Number and proportion of cells in each cluster

Because the DOL cluster was unique to the striatum ipsilateral to MCAO, we assessed differential gene expression and gene ontology (GO) of biologic processes in the DOL cluster compared to the nearest neighbor clusters on the dendrogram: Pre-OL2, MFOL and MOL (figure 5). The DOL cluster was relatively more enriched in GO terms relating to cellular responses to stress and antigen processing and presentation, among others (figure 5A). In contrast, GO terms relatively suppressed in the DOL cluster compared to neighbor clusters included biologic processes involved in synthesis and metabolism of myelin components, as well as clearance of amyloid beta (figure 5B). The top 50 genes up- and down-regulated in the DOL cluster compared to neighboring clusters are shown in supplemental tables 3 and 4 respectively.

**Figure 5:**
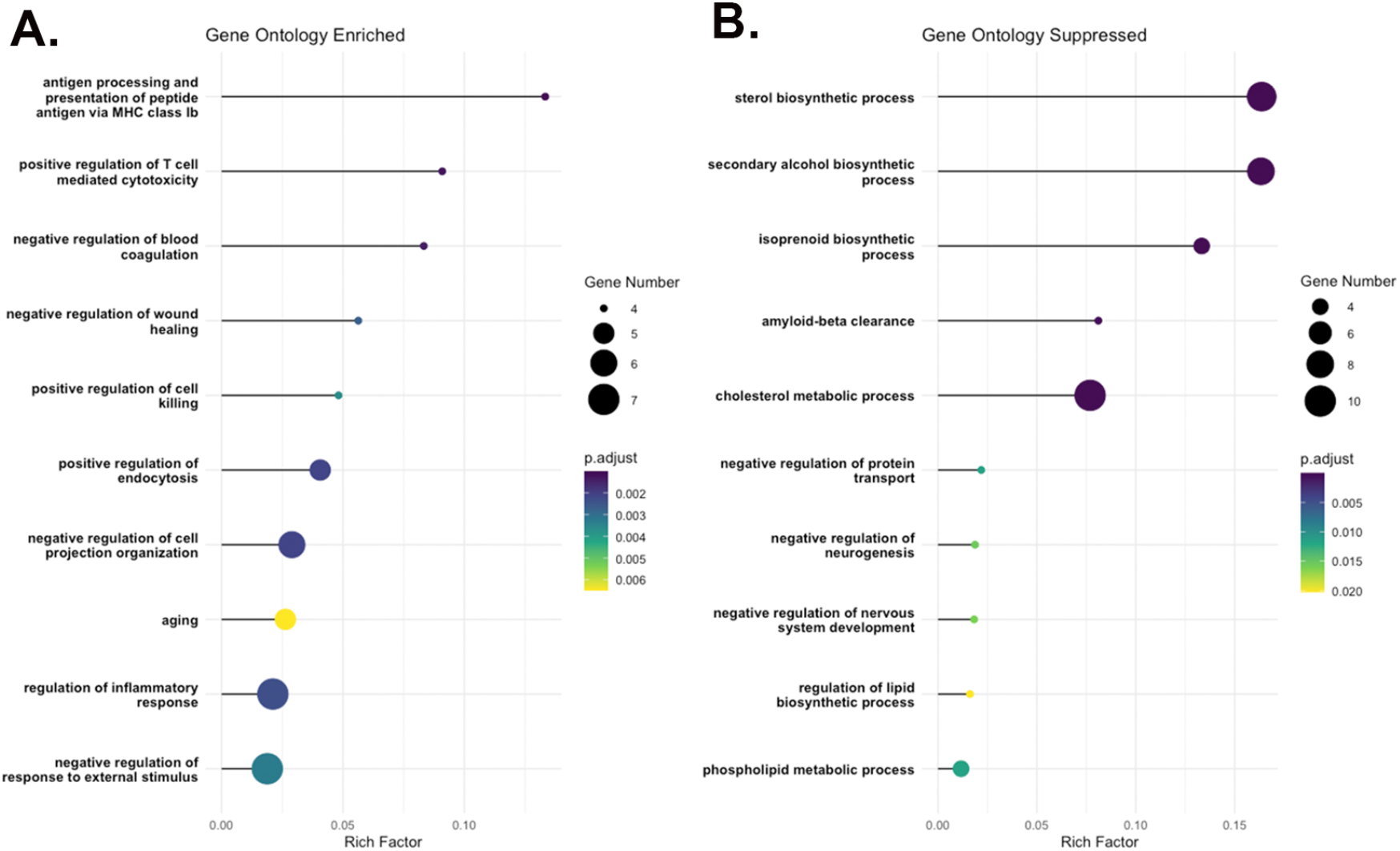
Gene ontology enrichment analysis of DEGs in the DOL cluster compared to nearest neighbors on the phylogenetic tree (Pre-OL2, MFOL, MOL). The Rich Factor is the ratio of differentially expressed gene numbers annotated in each biologic pathway term to all gene numbers annotated in the pathway term. A higher Rich Factor indicates a greater degree of pathway enrichment (A) or suppression (B) in the reactive cell cluster.

We next examined which upregulated genes defined the DOL cluster. Figure 6A shows differential expression of genes specifically upregulated in the DOL cluster. This includes many immune response genes, *IL33, C4b*, and *Klk6*, which is thought to promote inflammation via cleavage of protease-activated receptor 1 and 2[37], and *Serpin3N*, which encodes anti-chymotrypsin and is known to be upregulated in neuroinflammation[38]. Other upregulated genes specific to the DOL cluster were the long non-coding RNA *Neat1*, and *Jph4*, which encodes junctophilin 4, a protein responsible for tethering surface ion channels to the endoplasmic reticulum in excitable cells[39]. The fold change in gene expression in the DOL cluster compared to each other cluster of these reactive-defining markers is listed in supplemental table 5. Figure 6B shows genes that are over-expressed in the DOL cluster, but which are also expressed at varying levels in other clusters. These include a diverse set of genes that can be grouped into those involved in MHCI expression, neurotransmitter homeostasis and ion channel components, extracellular signaling, and response to oxidative stress. Table 2 lists descriptions of genes depicted in figure 6B. Fold change in gene expression in the DOL cluster relative to each other clusters of this gene set is listed in supplemental table 6.

**Table 2:** genes differentially upregulated in the DOL cluster.

| MHC I genes |  |
| --- | --- |
| B2M | Encodes beta 2 microtubulin, a component of MHC I. |
| H2-D1 | Encodes histocompatibility 2, D region locus1. Important for endogenous antigen presentation by MHC I via the endoplasmic reticulum. |
| Neurotransmitter homeostasis and ion channel components. |  |
| Glul | Encodes glutamine synthetase. Important for glutamate turnover and maintaining excitatory transmission. Has been shown to be expressed in oligodendrocytes in caudal mouse brain and spinal cord[40]. |
| Car2 | Encodes carbonic anhydrase 2. Carbonic anhydrases regulate intra- and extra-cellular pH transients and therefore influence proton-sensitive membrane receptors such as GABA and NMDA receptors[41]. |
| Hcn2 | Encodes hyperpolarization-activated cyclic nucleotide gated ion channels. HCN's contribute to neuronal excitability, but have also been found to regulate myelin sheet length and help maintain oligodendrocyte resting membrane potential[42]. |
| Atp1b3 | Encodes an isoform of the Na-K ATPase pump beta subunit. In naive adult mouse brain, the b3 isoform has low expression compared to the b1 and b2 isoforms[43]. |
| Extracellular signaling |  |
| Ptgds | Encodes prostaglandin D2 synthase. Prostaglandin D2 has a variety of neuro-modulatory roles within the CNS[44-48]. |
| ApoE | Encodes apolipoprotein E (ApoE). ApoE peptides are secreted from cells in lipidated aggregates and bind to a variety of cell types including neurons, microglia, and endothelial cells. Extracellular ApoE is important for modulating inflammation, clearing cellular debris, and promoting neurite outgrowth and dendritic spine formation[49]. |
| Sec11c | Encodes a catalytic subunit of the serum peptidase complex, which is important for generation and processing of excreted proteins. The serum peptidase complex removes signaling peptides from pre-proteins as they are translocated to the ER from ribosomes[50]. |
| Oxidative stress response |  |
| Gstp1 | Encodes glutathione-S-transferase pi (Gst-pi), which catalyzes a number of detoxification reactions within cells. Gst-pi in the brain is specific to oligodendrocytes and is upregulated in response to oxidative stress. Gst-pi inactivates harmful by-products of oxidative stress[10, 51, 52]. |

**Figure 6:**
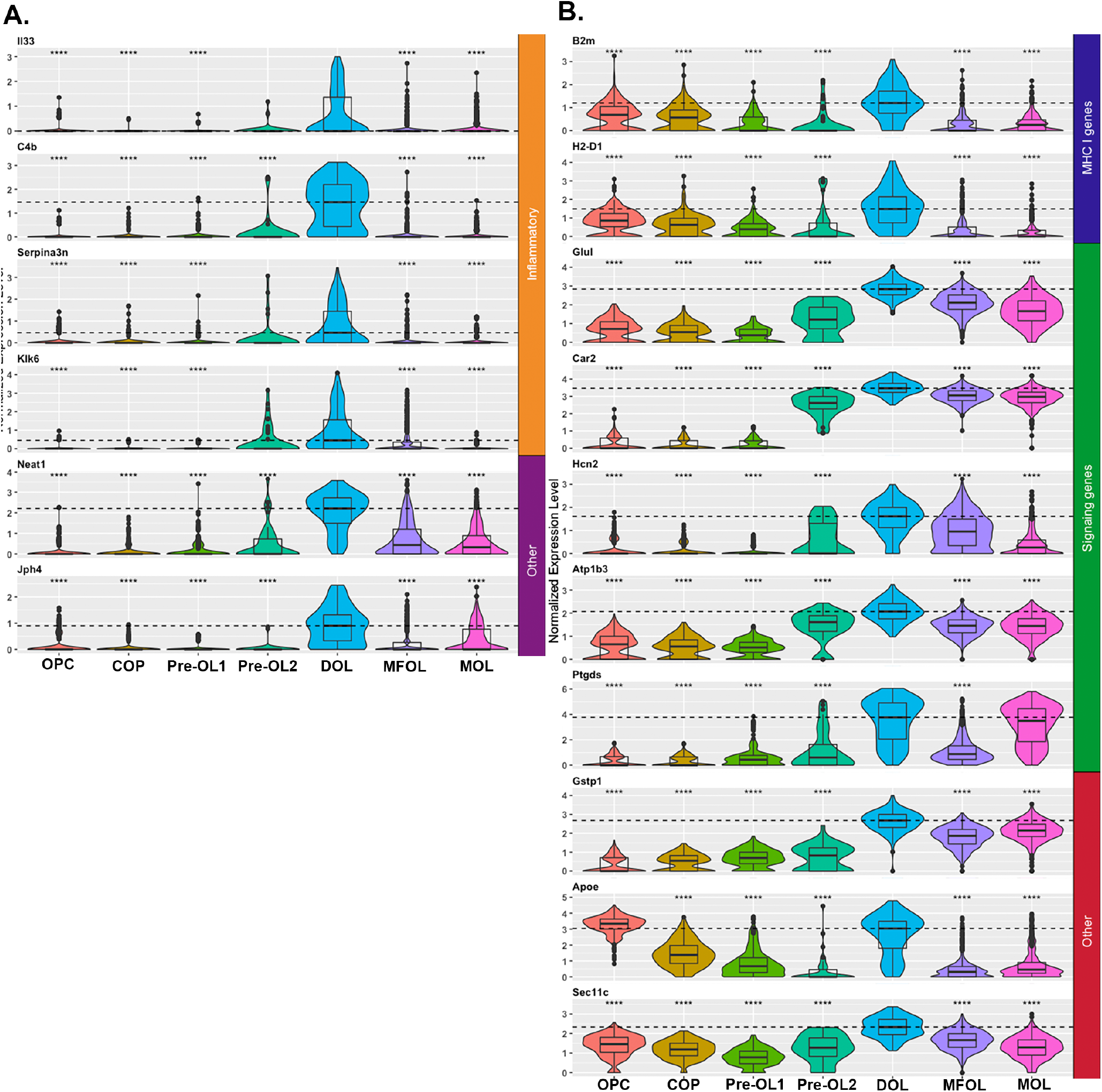
Gene expression in the DOL Cluster. **A**: Normalized gene expression of markers that define the DOL cluster.**B**: Gene expression of markers increased in the DOL cluster, but also expressed in other clusters. Boxplots demonstrate median and IQR of normalized gene expression levels. Dashed line marks median normalized gene expression in the reactive cluster. For each cell cluster DE analysis was preformed compared to the reactive cluster. The adjusted p-value for the average log2 fold change in gene expression between the DOL cluster and each other cluster was calculated using the Wilcoxon rank sum test with correction for multiple comparisons. **** denotes significant decrease in average log2FC compared to reactive cluster with adjusted p-val<0.0001.

We next compared differential gene expression between oligodendrocyte cells from ipsilateral versus contralateral striatum in each cell cluster (figure 7A). Relatively few genes were differentially regulated in the ipsilateral OPC and COP clusters. The MFOL cluster had the most differentially upregulated genes, and the MOL cluster had the most differentially downregulated genes. Ipsilateral cells in the MFOL and MOL clusters overexpressed the interferon early response genes *Ifi27* and *Ifi27l2a*. In addition, several genes overexpressed in the reactive cluster were also overexpressed in ipsilateral cells in the MFOL/MOL cluster, including the MHCI transcripts *B2m* and *H2-D1*, as well as *Gstp1, Neat1, Apoe and Ptgds*. Interestingly, the *Bcas1* gene was over-expressed in ipsilateral MFOL and MOL clusters relative to contralateral. *Bcas1* expression is thought to define newly differentiated myelinating cells in the adult brain[34]. Both *Socs3* and *Junb* were over-expressed throughout the lineage. Many fewer genes are down-regulated in ipsilateral cell clusters (figure 7b). Many of the genes down-regulated in ipsilateral cells clusters are genes for ribosomal proteins and genes involved in post-translational RNA modification. Interestingly the myelin associated gene *Mbp* was upregulated in the ipsilateral MOL cluster, but *Mobp* was down-regulated in the MFOL cluster. Otherwise, myelin associated genes were not significantly differentially regulated between ipsilateral and contralateral clusters. Genes upregulated and downregulated in ipsilateral hemisphere compared to contralateral in each cluster are listed in supplemental tables 7 and 8 respectively.

**Figure 7:**
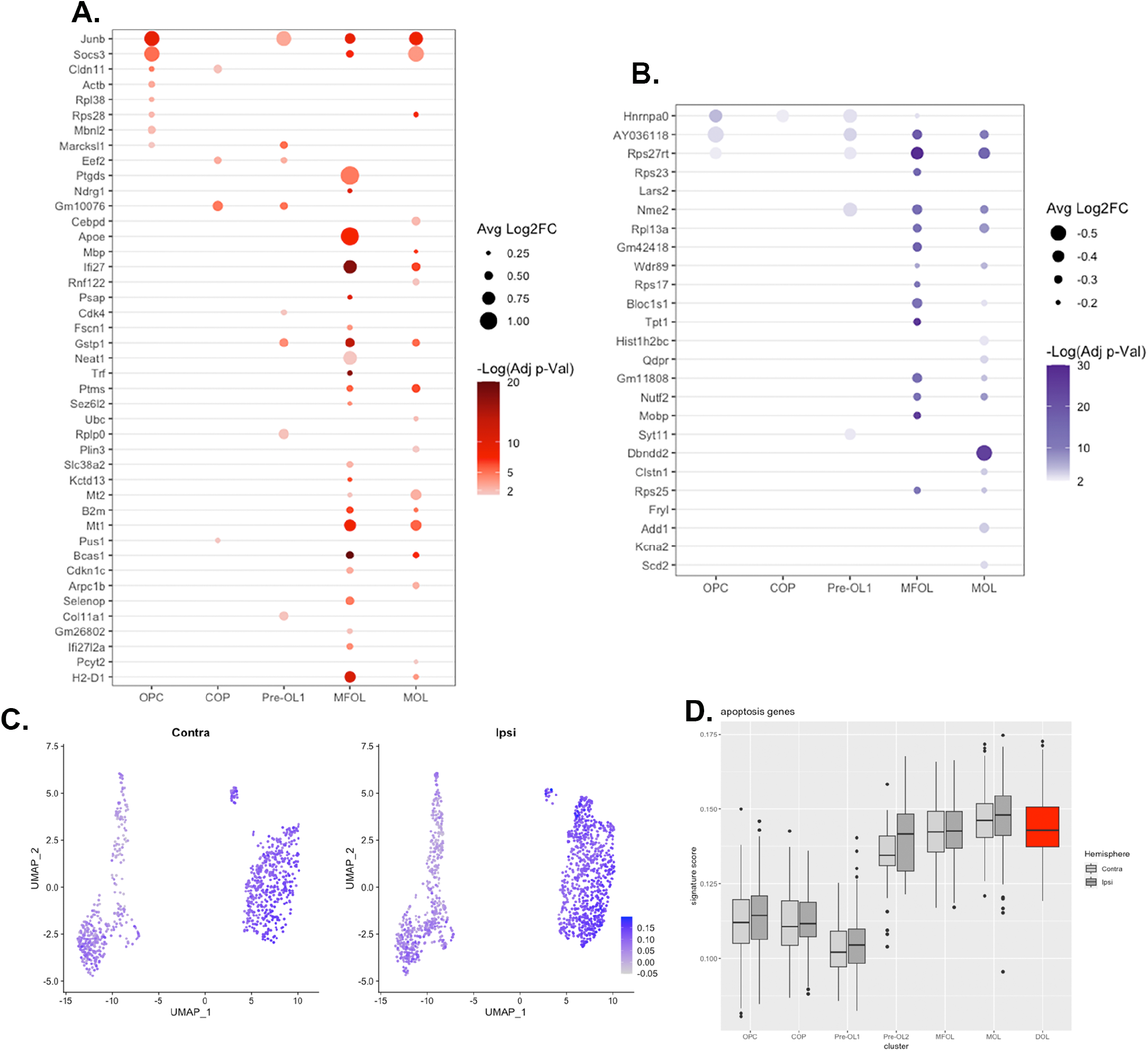
Differentially expressed genes in ipsilateral striatum oligodendrocytes in each cluster. **A:** Significantly upregulated genes in OLs from ipsilateral striatum compared to contralateral in each cluster. **B:** Significantly downregulated genes in ipsilateral OLs compared to contralateral from each cluster. There were no differentially expressed genes between contralateral and ipsilateral hemispheres in the Pre-OL2 cluster (excluded from figure). The Log base 2 of fold change is shown, so that an avg log2FC of 1 is a doubling in gene expression. Only genes with a log2FC> 0.25 are shown. The adjusted p-value for the average log2FC in gene expression between contralateral and ipsilateral cells in each cluster was calculated using the Wilcoxon rank sum test with correction for multiple comparisons. The adjusted p-value is represented on a negative Log_10_ scale, such that 1.3 represents an adjusted p-value of 0.05. Only values with adjusted p-value<0.05 are shown. **C-D:** Apoptosis signature gene scores using the the GSEA Molecular Signature Database Hallmark Apoptosis gene set were assigned to each cell using the UCell package in R. Apoptosis scores for each cluster in ipsilateral and contralateral hemisphere are shown as a feature plot(C) and bar graph (D). In D, for the reactive cluster only ipsilateral is shown as there were only 4 cells in the reactive cluster in the contralateral group.

Because there was a decrease in CC1^-^ oligodendrocytes between 14-and 28-days post-MCAO without a clear increase in CC1^+^ cells over this time, we queried our scRNA-seq data for apoptosis-associated gene expression in contralateral and ipsilateral samples across cell clusters (figure 7C-D). Each cell was scored as previously described[53], based on the 161 genes in the GSEA Molecular Signature Database Hallmark Apoptosis gene set (https://www.gsea-msigdb.org/gsea/msigdb/human/geneset/HALLMARK_APOPTOSIS.html)[54]. The apoptosis gene score was overall higher in more mature clusters and the DOL cluster compared to immature clusters, but was not markedly higher in ipsilateral cells from any clusters (figure 7C-D).

We sought to validate whether DOLs as defined in our seq data set were identifiable in situ in the ipsilateral striatum in tissue sections 14 days after p10 MCAO. Using fluorescent in-situ hybridization with probes for *Sox10* mRNA (to identify oligodendrocytes), paired with two probes for mRNA over-expressed in the reactive cluster in our seq data (*B2m*, and *Neat1*). We found that in striatum ipsilateral to MCAO, cells in which probes for *B2m* and *Neat1* colocalized with *Sox10* (figure 8A top row and figure 8B) were present. While each transcript was present in contralateral striatum, colocalization of transcripts was not present within the same cells (figure 8A bottom row). We performed IHC 14 days post-MCAO to determine if oligodendrocytes in ipsilateral striatum had increased expression of genes differentially expressed in our SEQ data (figure 8C&D). We found that both Gst-pi^+^ cells and MHCI expression were increased in ipsilateral striatum (figure 8C bottom row) compared to contralateral (figure 8C top row). Gst-pi expression in the brain is specific to oligodendrocytes[10, 52]. Most MHC-I reactivity co-localized with microglia (Iba-1^+^), which are in close contact with Gst-pi^+^ cells (figure 8C bottom row). However, high-power 3d reconstruction demonstrated that MHC-I is also present within the cell body of GST-pi^+^ cells, and on the cell surface (figure 8D-G).

**Figure 8:**
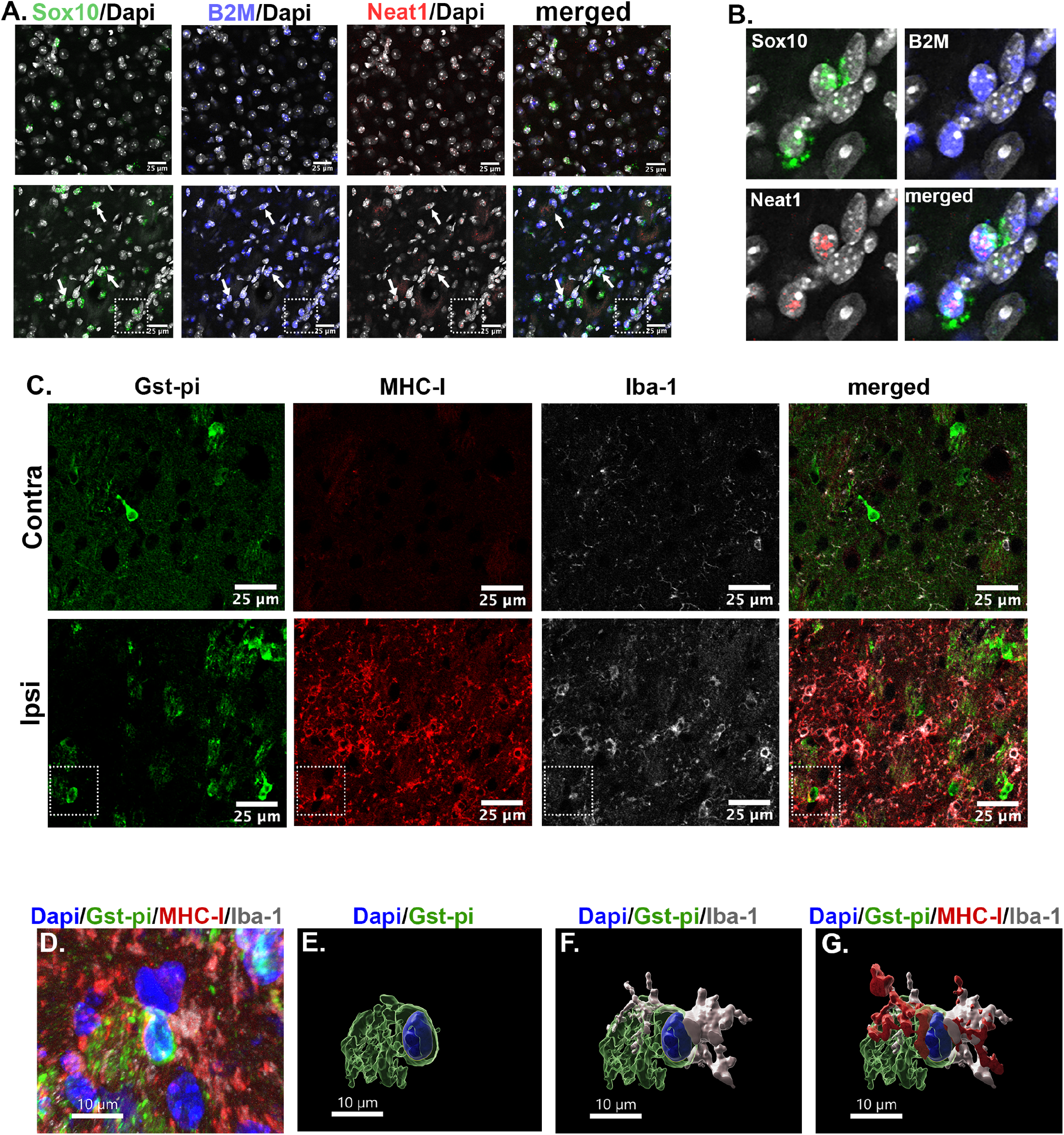
IHC and FISH show expression of DOL cluster genes in ipsilateral striatum. **A-B**: RNA-scope FISH of contralateral and ipsilateral striatum 14 days after MCAO with probes for Sox10, B2M and Neat-1 mRNA. Nuclei were labelled with Dapi (blue). In Contralateral striatum (top row) cells expressing all 3 mRNA were not identified. There were several cells that expressed all 3 mRNA in ipsilateral striatum (bottom row, arrows and white box). **B:** Magnification of white box in A. **C:** IHC for Gst-pi, MHC-I, and Iba-1 in contralateral (top row) and ipsilateral (F-I) lateral striatum 14 days post-MCAO. Both Gst-pi and MHC-I expression are increased in the ipsilateral striatum. In Ipsilateral striatum Most MHC-I co-localizes with Iba^+^ microglia, but there are Gst-pi^+^ cells expressing MHC-I (arrows and dashed box). **D:** High power view of area in white box in C. **E-G**: 3D reconstruction of GSTpi+ cell in D using Imaris software.

Recent data has suggested that in inflammatory demyelination, OPC’s respond to interferons (IFN) by increasing expression of MHCI and MHCII genes as well as genes involved in antigen processing and presentation[16]. To determine whether a similar phenomenon was occurring after neonatal stroke, we examined gene expression of interferon early response genes as well as MHCI and II genes in our oligodendrocyte clusters (figure 9A). We generated a single cell composite score for each set of genes in each cluster from ipsilateral and contralateral hemisphere using previously published methods[53] (figure 9B). In most cell clusters composite expression of IFN response genes and MHCI genes is increased in cells from the ipsilateral striatum compared to contralateral, as well as increased in the DOL cluster (figure 9B&C). However, MHCII genes are also not significantly expressed in clusters from either hemisphere (figure 9D). Kenigsbuch et al have recently described disease associated oligodendrocytes across different mouse models of dementia and experimental auto-immune encephalitis (EAE) that share a common gene expression signature [1]. We evaluated expression of this DOL signature in our data set (figure 9E-F). There was a large overlap in expression of genes identified by Kenigsbuch et al as defining DOLs and in the DOL cluster in this study. The composite gene score of genes listed in figure 9E was much higher in the DOL cluster than in other clusters of either hemisphere (figure 9F).

**Figure 9:**
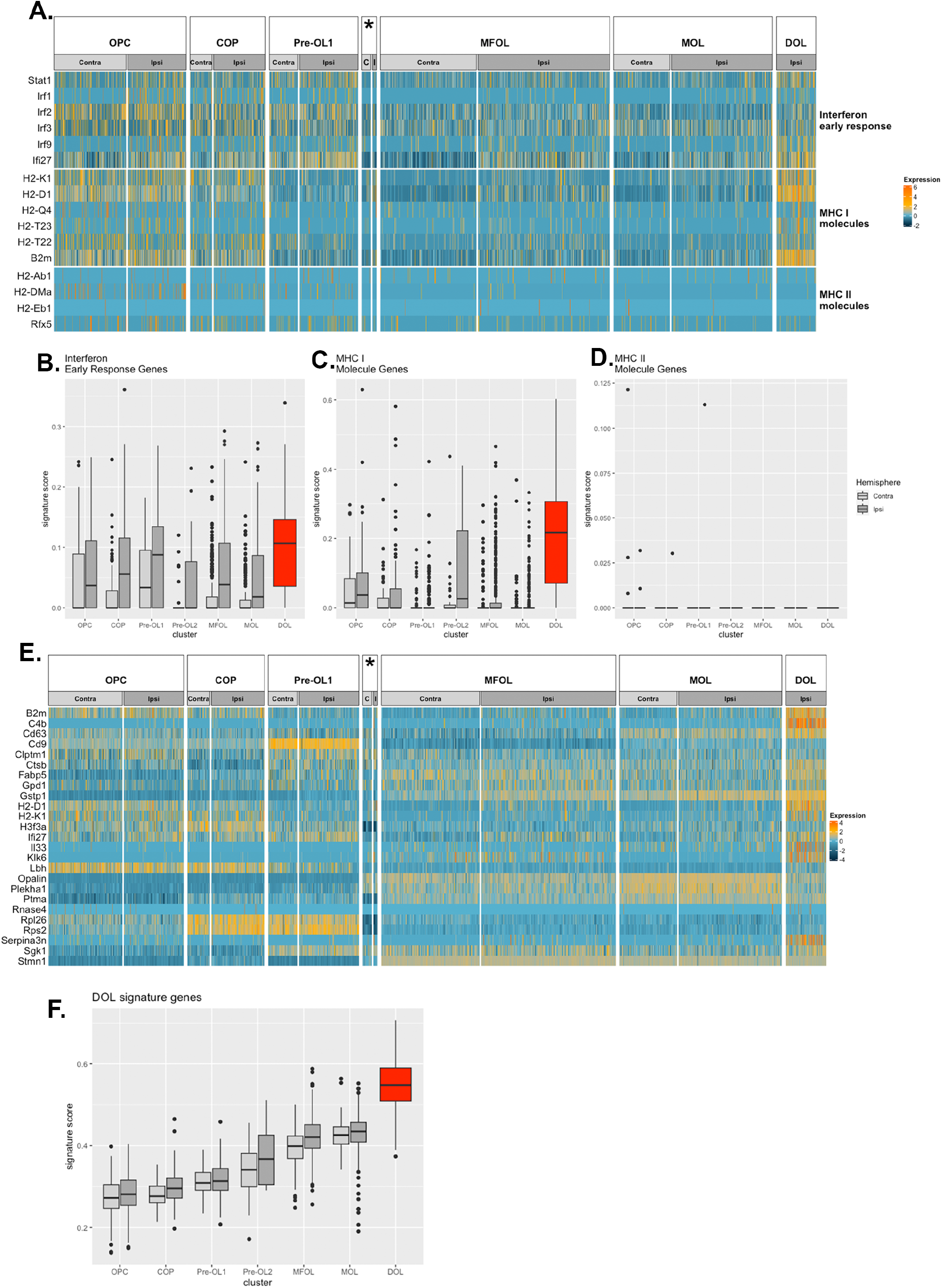
IFNy response and antigen presentation gene expression. **A**. Heat map of gene expression in cells from ipsilateral and contralateral striatum in each cluster and in the DOL. **B-D:** Signature gene scores for each category were calculated using the Ucell package for R. **E:** Heat map showing gene expression of genes associated with a DOL population described in models of AD and EAE[1]. **F:** The genes in E were used to generate a signature gene score for cells in each cluster.

## DISCUSSION

In this study we explored changes in white matter and oligodendrocyte populations in a neonatal model of arterial ischemic stroke. We found a histologic increase in oligodendrocyte density in the white-matter rich lateral striatum two weeks after MCAO. Our single cell RNA sequencing data provided a more granular insight in to changes in oligodendrocyte sub-populations at this time point. The most surprising finding in this study was the existence in ipsilateral striatum of a subset of “reactive” DOLs that were nearly non-existent in the contralateral hemisphere. This DOL cell cluster appeared to be closer in maturation to early myelinating oligodendrocytes than to less mature stages. In prior studies using the Rice-Vannucci model of hypoxia and ischemia (HI), Back et al described a sub-set of pre-myelinating (O4^+^O1^+^) oligodendrocytes that differentiate after HI with a “reactive” morphology[9]. The production of these “reactive” cells is greater after HI at term-equivalent compared to earlier in development, suggesting that their generation may be related to the relative abundance of pre-myelinating OL’s at this age. Beyond a morphologic description of these cells, they have not been investigated in the setting of perinatal brain injury. The DOL cluster in this study expressed a number of genes suggesting an immune/inflammatory profile, including upregulation of MHCI genes, *Il33* and compliment genes. Disease-specific oligodendrocytes have now been recognized in a number models of white matter injury in adult rodents. Falcão et al found disease-specific mature oligodendrocytes in mouse experimental autoimmune encephalitis (EAE), which share transcriptional similarities to the post-stroke reactive cells in this study, including expression of *Serpina3n* and MHC-1 genes[14]. Similarly, a “reactive” cluster of oligodendrocytes was identified in an Alzheimer’s disease (AD) mouse model, defined by *Serpina3n, C4b*, and *H2-D1*[55], all genes upregulated in the reactive oligodendrocyte cluster after neonatal stroke in this study. Recently, RNAseq data was integrated from EAE and AD mouse models, demonstrating that DOLs across these models have a shared gene expression signature, which overlaps with the DOL expression pattern in our model of neonatal stroke. As far as we are aware, this is the first study reporting immune oligodendrocytes in ischemic stroke or in a perinatal brain injury model.

The signaling pathways that give rise to immunogenic oligodendrocytes are unclear. We found that interferon early response genes, such as *Ifi27* were upregulated in the reactive cluster, as well as in the ipsilateral cells in the MFOL and MOL cluster. IFN has long been known to induce MHCI expression in oligodendrocytes throughout the lineage[16, 56]. We found that MHCI genes *B2m* and *H2-D1* were differentially expressed in the more mature MFOL/MOL clusters ipsilateral to MCAO and in the reactive cluster, but not in OPCs. We also did not see differential expression of MHCII genes in any clusters, suggesting that the immune response to IFN and other signals in the context of stroke and early development is distinct from other disease models where upregulation of MHCII on oligodendrocytes has been described[14]. IFNγ is not increased in the acute time period after p7 MCAO in rats[57]. IFNγ is largely thought to be produced by lymphoid cells, although microglia *in vitro* can produce IFNγ under the influence of IL-12 and IL-18[58]. One reason for the paucity of IFNγ acutely after neonatal stroke is likely that the blood brain barrier is relatively preserved acutely after neonatal mouse MCAO, preventing lymphocyte infiltration[59]. However, there is vessel degeneration and dysfunctional angiogenesis chronically after neonatal stroke, suggesting that lymphocyte infiltration may occur at delayed timepoints[60]. Reactive OL generation in our model of neonatal stroke may be due to immune cell signals other than lymphocytes. Zhou et al demonstrated that the presence of reactive OLs expressing *C4b* and *Serpina3n* in AD phenotype mice was shown to be at least partially dependent on microglial activation[55]. We did not investigate lymphocyte infiltration or quantify microglial activation in this study, but did observe increased microglial MHC-I expression in injured striatum, suggesting a chronically activated phenotype.

In our hands 60 minutes of MCAO in p10 mice produces significant neuronal death in lateral striatum by 3 days post-injury. While there was some loss of MBP^+^ myelin in the same area, the radial white matter fibers in the area of maximal neuronal loss were overall relatively preserved. This suggests that, contrary to rodent models of ischemic injury in preterm-equivalent animals, white matter is relatively resistant to acute injury in term-equivalent animals. In humans, myelination begins shortly after the onset of immature oligodendrocyte expansion, at around 30 weeks gestational age (GA), although this varies by brain region from about 20-33 weeks GA[61, 62], and continues into early adulthood with peak myelination in late infancy/early childhood[63-65]. Thus, at the time of stroke in full term neonates myelination is underway but is not at its most active point. In mice, this is generally agreed to correspond to p10, with p20-21 corresponding to peak myelination[66]. Ahrendsen et al previously found when MCAO is performed at p20-25, just after the time of peak myelination, striatal white matter is similarly preserved ipsilateral to the injury, with much greater white matter destruction in adult animals (8-12 weeks old) when active myelination has slowed[10].

Prior research suggests that the maturational state of white matter determines its relative vulnerability to ischemic injury. In post-mortem human brains the oligodendrocyte population is primarily made up of early and late oligodendrocyte progenitor cells prior to 28-weeks GA in peri-ventricular white matter, with rapid expansion of early myelinating immature oligodendrocytes (O4^+^O1^+^) between 28- and 41-weeks GA[61]. The expansion of immature oligodendrocytes corresponds to the end of the most vulnerable time period for periventricular leukomalacia, the predominant ischemic white matter injury of prematurity. In rodents, P2-5 corresponds to a similarly vulnerable time period for white matter injury, in which little myelination exists and oligodendrocytes consist of primarily pre-myelinating progenitors, but by p7 immature oligodendrocytes predominate[67]. Back et al have shown that regardless of the timing of ischemic injury, late oligodendrocyte precursors are specifically vulnerable to ischemic injury, with earlier progenitors and immature early myelinating oligodendrocytes showing resistance to death after ischemia[9]. Therefore, the relative vulnerability or resistance of white matter to injury appears to be due to the makeup of the oligodendrocyte lineage at the time of injury. Resistance to ischemia during times of active myelination may be due to increased anti-oxidant capacity of myelinating oligodendrocytes compared to mature oligodendrocytes in adult animals[10]. Similarly, we found that two weeks after p10 MCAO, more mature oligodendrocytes from ipsilateral striatum had significantly increased expression of *Gstp1* mRNA. These data suggest that white matter vulnerability in the full-term neonatal period is more similar to that of childhood than to preterm infants. Furthermore, the resistance of white matter to ischemia during this time period may be due to the predominance of early myelinating oligodendrocytes with greater anti-oxidant capacity.

Although white matter in ipsilateral striatum is relatively preserved after p10 MCAO, the impact that the observed gene expression changes in reactive and other mature oligodendrocyte clusters have on long-term white matter recovery and myelination is unclear. We found a dramatic increase in the density of myelinated axons within pencil fibers between 14-days(p24) and 28-days(p38) post-MCAO in both contralateral and ipsilateral striatum, consistent with the fact that this is the time period of most active myelination in mice. However, there were significantly fewer myelinated axons in ipsilateral pencil fibers at 28-days post-MCAO, despite the fact that there was no difference between hemispheres at 14-days MCAO. This parallels findings in human infants who suffered neonatal stroke, in which white matter tracts ipsilateral and contralateral to the infarct do not show differences in integrity (measured by MRI diffusion tensor imaging) several weeks after stroke, but have deterioration up to 24 months after the stroke[8]. In addition, by 28-days post-MCAO myelin had more dysplastic features in ipsilateral white matter compared to contralateral. These findings may be due to reduced OL myelinating capacity. The reactive OL cluster in this study had reduced expression of genes involved in lipid biosynthesis compared to adjacent cell clusters, suggesting they are less involved in myelin production. Transgenic mouse models in which MHCI genes are over-expressed specifically in more mature oligodendrocytes have shown that this results in MHCI molecule accumulation in the endoplasmic reticulum (ER), leading to ER stress and hypomyelination[68, 69]. Further studies are needed to determine whether oligodendrocytes ipsilateral to ischemia have chronically reduced myelinating potential in the developing brain.

We observed a robust oligodendrocyte proliferation in the injured striatum at 3-7 days pot-MCAO by EdU labelling of dividing cells during this time. In addition, oligodendrocytes that were newly-born during the sub-acute injury period persisted as immature (CC1^-^) oligodendrocytes two weeks post-MCAO. However, we did not see a commensurate increase in mature EdU^+^ oligodendrocytes at 28 days post-MCAO, suggesting that the majority of newly-born oligodendrocytes after MCAO do not survive and mature in to myelinating cells. Oligodendrocyte proliferation after white matter injury is a long-observed phenomenom. Back et al found that NG2^+^ progenitors proliferated after hypoxic ischemic injury in rodents, at both preterm (p2) and term (p7) equivalence, within 24 hours of injury[9]. Proliferation of progenitors after ischemia is not limited to the young brain. Increased density of OPCs has been demonstrated 2-weeks post-reperfusion in the peri-infarct area in adult rodents[70]. Ahrendsen et al previously found that after stroke in juvenile mice (p21-24) there is early OPC proliferation at 3-7 days post-MCAO, but in adult mice there is an initial loss of OPCs, followed by a rebound at 7-days post-MCAO[10]. However, the long-term fate of newborn OPCs after ischemia has not been well-studied. In the perinatal period, ischemia has been shown to inhibit OPC differentiation[71]. Extensive research in rodent models of *pre-term* brain injury suggest that tissue-environment factors inhibiting OPC differentiation include increased inflammatory cytokines, reactive oxygen species, and other factors produced by activated microglia and astrocytes. However, factors that influence OPCs maturation, or lack thereof, in ischemic injury at *term* equivalent have not been widely explored.

Single cell clustering and differential gene expression in our data set support our histologic findings. In the cells isolated from ipsilateral striatum there was a significant *decrease* in the OPC population, and a significant *increase* in the next most mature sub-population (COP) 2 weeks post-MCAO. This could indicate that OPCs were selectively lost with compensatory proliferation of COPs. Alternatively, OPCs may have been driven to differentiate into COPs, depleting the OPC pool, with subsequent proliferation and maturational arrest of the COP population. Differential gene expression in each cluster suggest the latter. The OPC cluster from ipsilateral striatum had significantly higher expression of the genes *Junb, Socs3, Actb*, and *Cldn11. Junb* encodes a subunit of the transcription factor AP-1, and is generally thought to antagonize cell proliferation [72]. *Socs3* generally provides negative feedback of cytokine-induced Stat signaling, thus inhibiting downstream pro-inflammatory gene expression[73]. Studies in models of spinal cord injury suggest that *Socs3* inhibits OPC proliferation, and may promote differentiation by inhibiting *Stat*[74]. *Actb* encodes Activin B, an arm of the TGFb super-family of ligands, and, has been shown to promote OPC differentiation in primary cell culture [75]. While ipsilateral expression of *Cldn11* is increased in the OPC cluster, the fold-change in expression compared to contralateral is even greater in the COP cluster. Claudin-11 is a tight-junction protein that forms a complex with oligodendrocyte specific protein (OSP) and promotes OPC proliferation[76]. Furthermore, in the COP cluster there is increased expression of pro-proliferation genes *Eef2* and *Cdk4. Eef2* encodes Eukaryotic elongation factor-2 (EF-2). EF-2 protein function depends on its phosphorylation status. Increased unphosphorylated EF-2 is leads to increased translation and cell proliferation[77]. *Cdk4* encodes Cyclin-D Kinase, which phosphorylates Cyclin-D, resulting in down-stream inactivation of Rb, transcription and cell cycle entry[78]. Together, upregulation of genes that promote differentiation in the OPC cluster and of genes promoting cell proliferation in the COP cluster suggest that early OPC progenitors differentiated into later progenitors (COPs), and that maturational arrest is occurring at this time. However, a limitation of our study is that scRNA Seq was only performed at one timepoint. Therefore, we cannot rule out that COPs progress to more mature stages after the 2-week post-MCAO timepoint and then undergo cell death. We did see increased expression of apoptosis-associated genes in more mature cell clusters at 14-days post-MCAO, but expression was similar between ipsilateral and contralateral samples. The local factors that result in OPC proliferation and maturational arrest/ maturation and apoptosis were not determined in this study. As previously discussed, Interferon early response genes were up regulated in more mature clusters from ipsilateral striatum, but not in COP or OPC clusters. However, *Socs3* was increased in several cell clusters ipsilateral to MCAO, including OPC. *Socs3* is upregulated in response to IFNγ - induced Jak/Stat signaling[79] and has been shown *in vitro* to reduce oligodendrocyte progenitor differentiation[80]. It is also possible paracrine factors produced by more mature oligodendrocytes in the ipsilateral striatum influence progenitor cells. The reactive cell cluster over expresses *Klk6*, which has been shown to limit OPC differentiation into myelinating cells[81]. On the other hand, reactive cells and ipsilateral MFOL/MOL cells have significantly upregulated expression of *Apoe*, and ApoE is known to promote oligodendrocyte differentiation[82].

There are several limitations of this study. The experiments were performed in male postnatal mice only. Although we are inducing focal ischemia at a time period before the classic sexual dimorphism seen in adult stroke models, there is growing evidence that sex differences occur in other models of neonatal ischemia[83-85]. Furthermore, the chronic time points that we studied overlap with the onset of sexual maturation (14 days post-MCAO, postnatal day 24-25), and into young adulthood (28-30 days post-MCAO, postnatal day 38-39). Therefore, future investigations into sex differences of oligodendrocyte and myelin changes after neonatal stroke are warranted. Another limitation of this study is that chronic morphologic and transcriptomic changes in microglia/macrophages, astrocytes, and leukocytes were not addressed. There is a large body of research addressing neuroinflammation after ischemia in adult and neonatal rodents, and was beyond the scope of this study. However, it is likely that the chronic phenotypic and functional changes in oligodendrocytes are influenced by other glia and immune cells, and is a compelling line of future study.

In summary, we show that oligodendrocyte proliferation and differentiation are significantly altered at chronic time-points after neonatal stroke, at a time when developmental myelination is maximal. Remarkably, more mature oligodendrocytes adopt an immune phenotype, which shares many gene expression similarities with immune oligodendrocytes in models of adult neurodegenerative diseases. This is the first study to show such a change in a stroke at any age and in a model of ischemic developmental brain injury. We propose that strategies to target oligodendrocyte behavior at sub-acute and chronic time-points after neonatal stroke may offer new ways to improve white matter repair and ultimately functional outcome.

---

**Supplemental figure 1:**
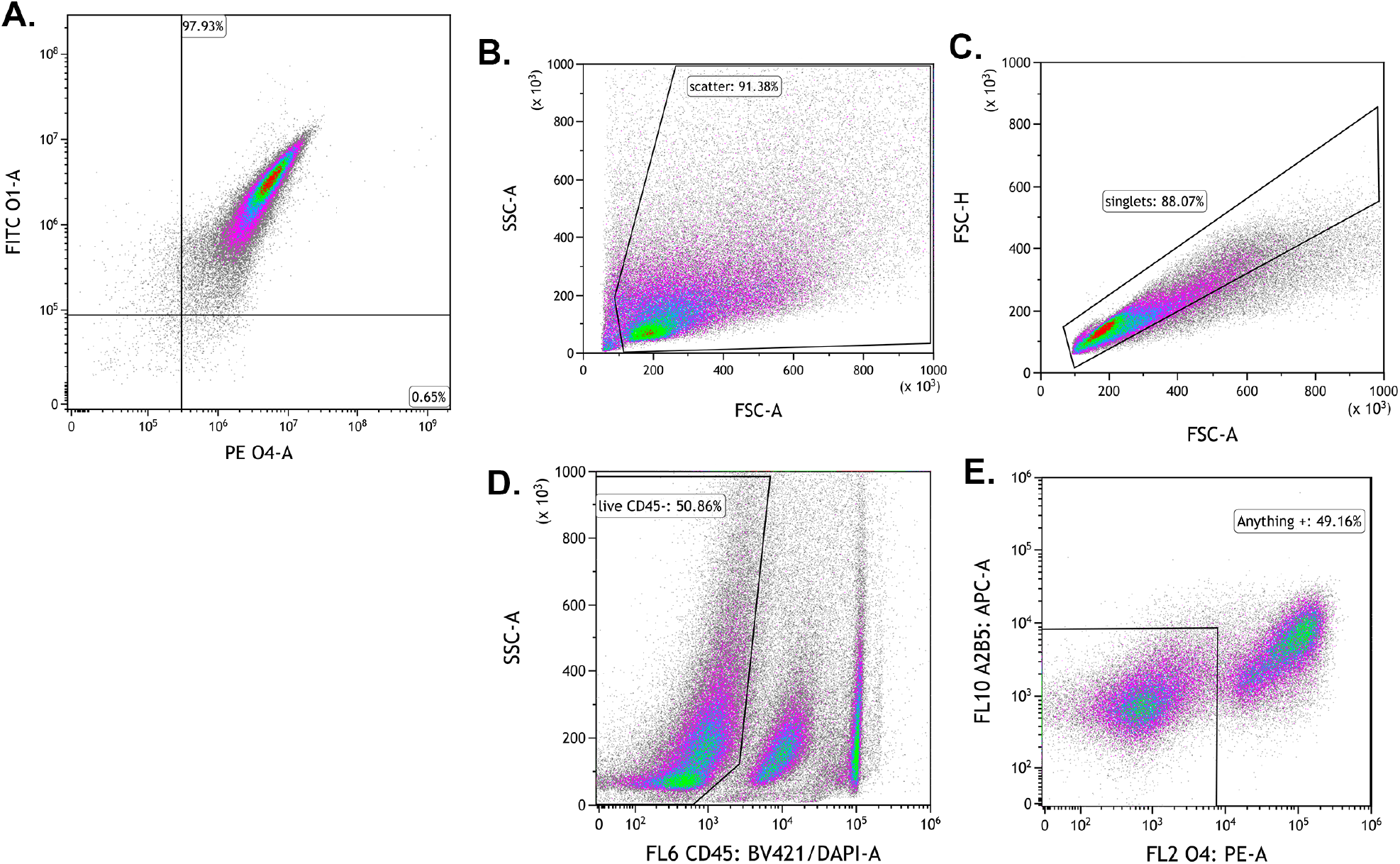
Oligodendrocyte marker selection and sorting strategy. **A:** Co-Expression of O1 and O4 surface markers. Nearly all cells positive for O1 were also positive for O4. **B-E:** Gating strategy for dissociated striatal cells. **B:** Debris was excluded using side scatter and forward scatter. **C:** Singlets were then selected. **D:** Cells negative for brilliant violet 421 (BV421) were then selected in order to exclude Dapi+ dying cells and CD45+ immune cells. **E:** Cells positive for the immature oligodendrocyte marker A2B5, or for the mature marker O4, and double expressing cells were sorted and collected for single cell sequencing.

**Supplemental figure 2:**
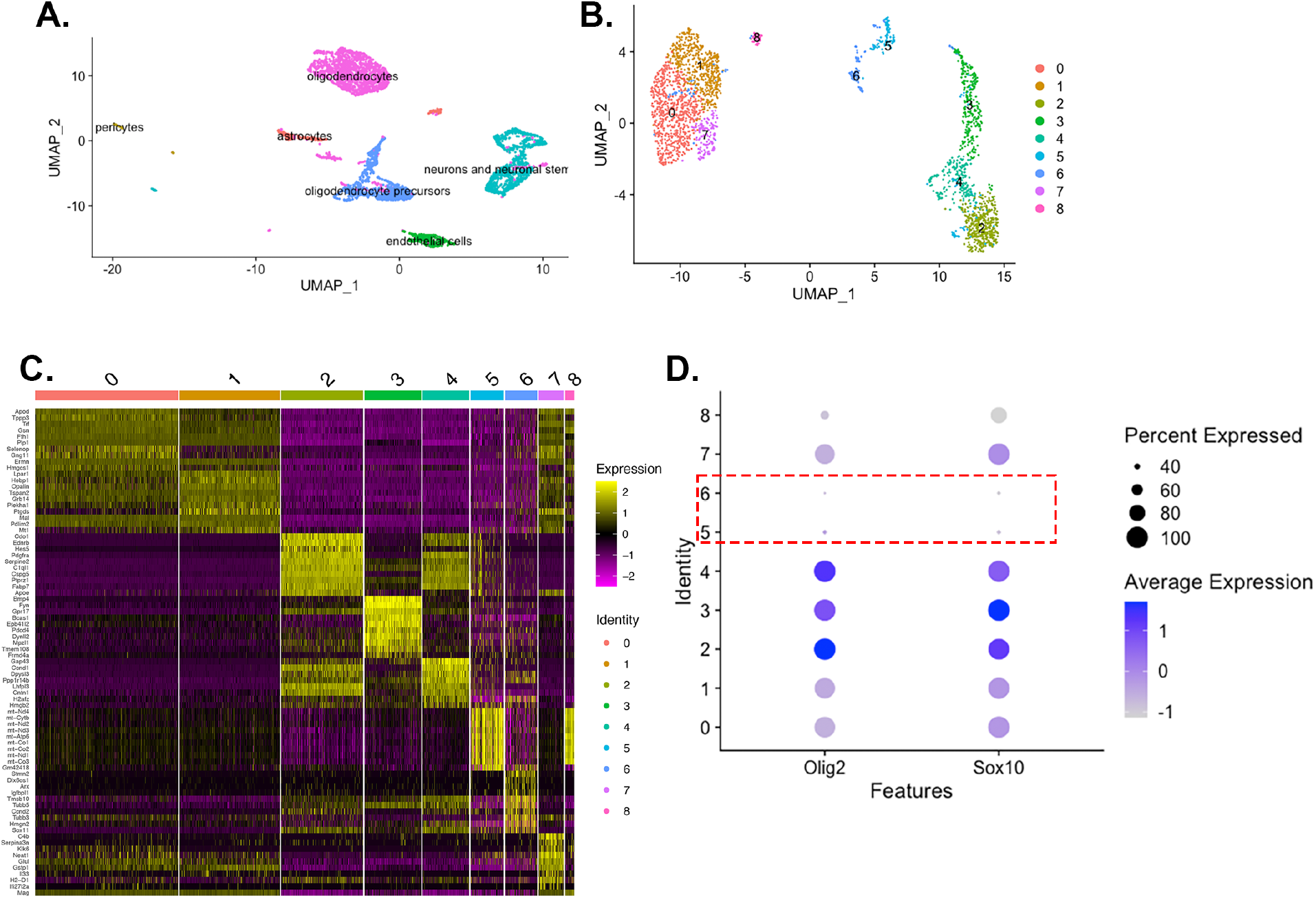
Determination of Oligodendrocyte Population. **A:** Initial cell clusters and cell identities based on the Tabula Muris gene expression database. **B:** Dimension plot of initial 9 clusters of oligodendrocyte lineage cells. **C:** Gene expression heat map of 9 initial oligodendrocyte lineage cell clusters. **D:** Olig2 and Sox10 relative expression in the 9 cell clusters from B. Clusters 5 and 6 were excluded based on low expression of the canonical oligodendrocyte genes Sox10 and Olig2.

**Supplemental figure 3:**
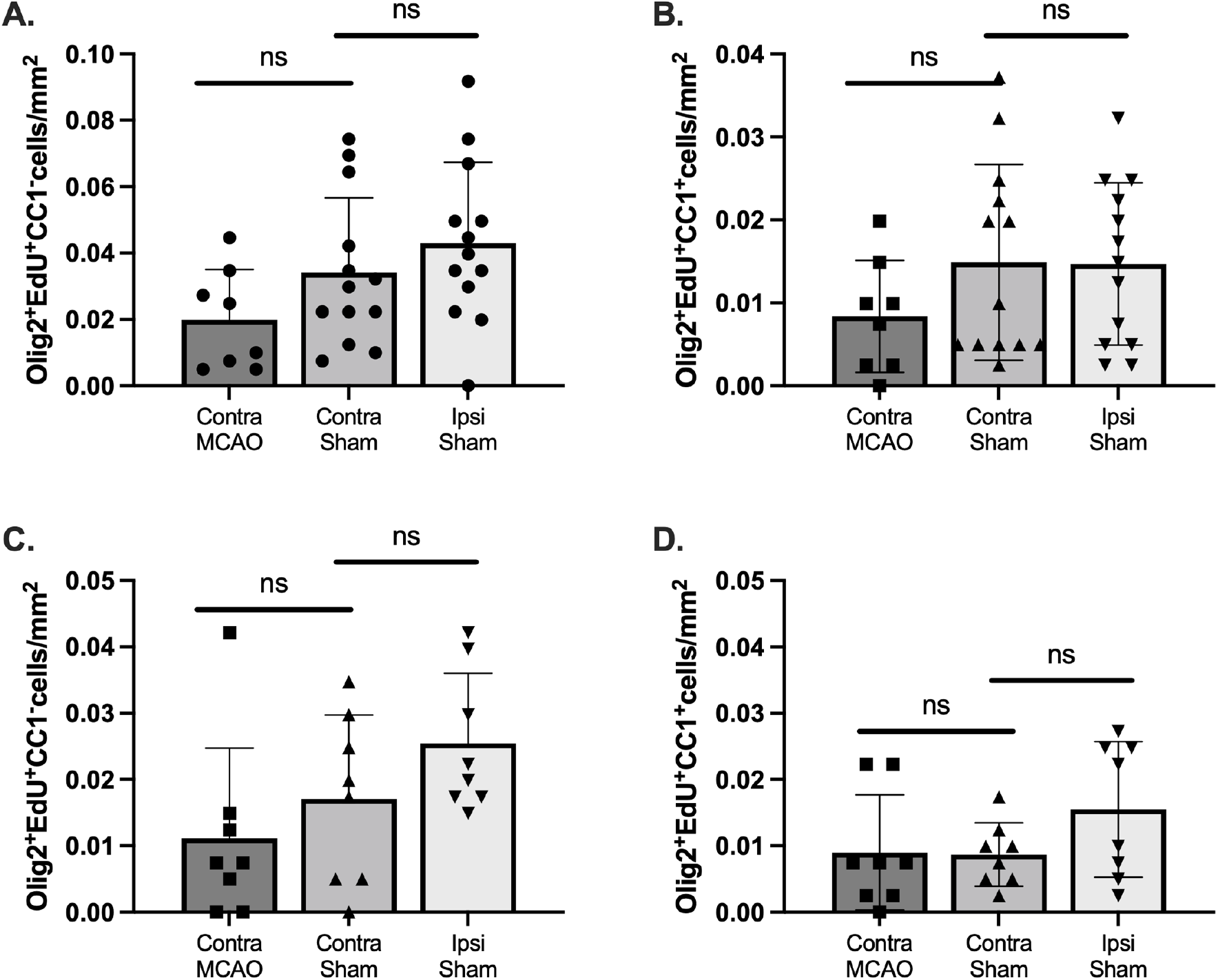
Density of immature and mature oligodendrocytes 14- and 28-days after MCAO. **A:** Density of immature (CC1-) oligodendrocytes (Olig2+) in contralateral hemisphere from MCAO-operated and both hemispheres from Sham-operated animals 14-days post-MCAO. **B:** Density of mature (CC1^+^) oligodendrocytes 14-days post-MCAO. **C:** Density of immature oligodendrocytes 28-days post-MCAO. **D:** Density of mature oligodendrocytes 28-days post-MCAO. A-C analyzed with Kruskal-Wallis test, D analyzed with 1-way ANOVA.

**Supplemental figure 4:**
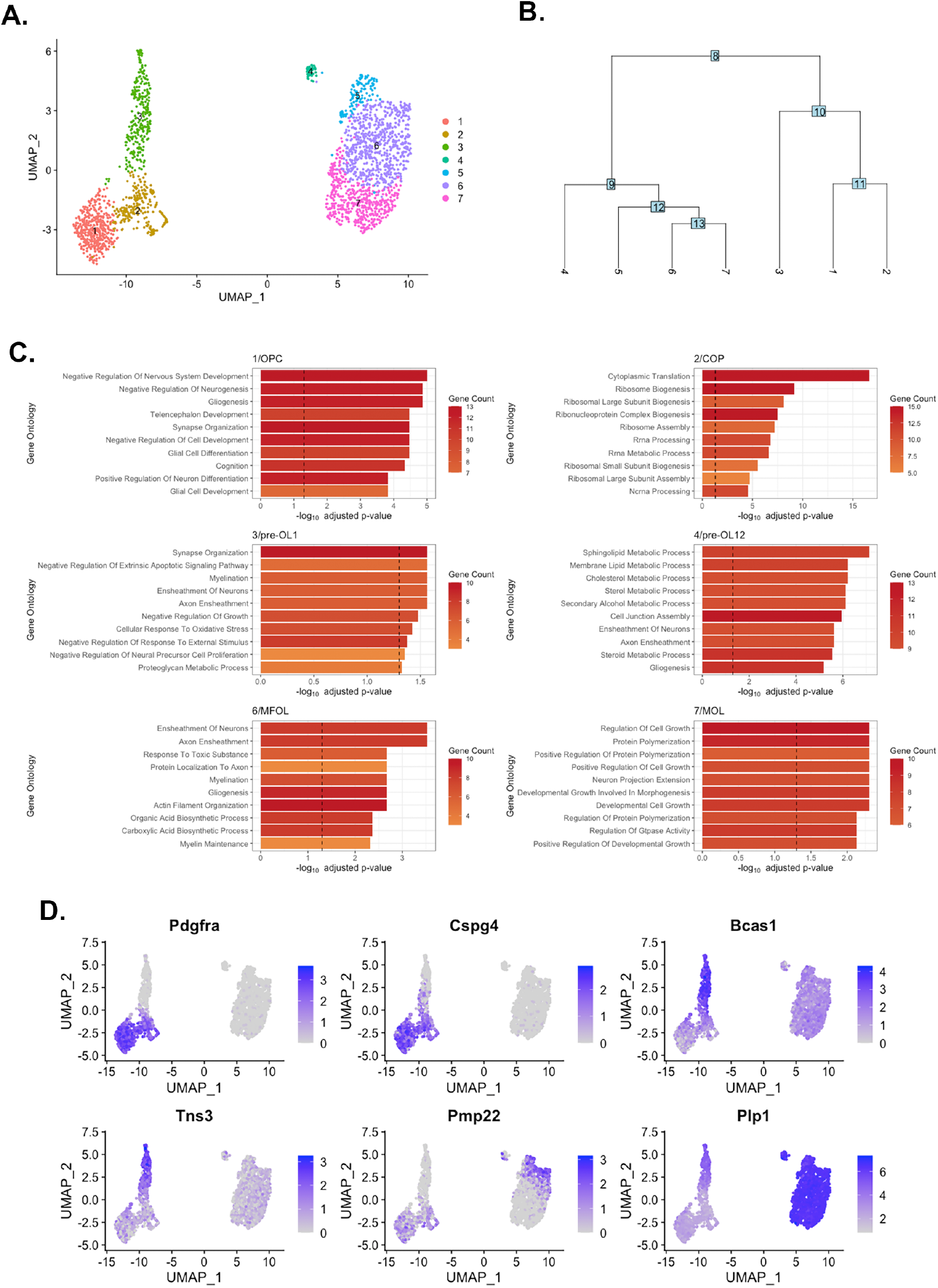
Oligodendrocyte cell cluster classification. **A:** Dimplot of final 7 oligodendrocyte clusters. **B:** Phylogenetic tree showing relationship between clusters. **C:** Gene ontogeny analysis of the top differentially expressed genes in each cluster. Each cluster graph is named with the original cluster number as well as the reassigned name based on analysis. **D:** Feature plot showing expression of OPC-specific genes (*Pdgfra* and *Cspg4*), Pre-myelinating oligodendrocyte specific genes (*Bcas1* and *Tns3*), and myelinating/mature oligodendrocyte specific genes (*Pmp22* and *Plp1*).

**Supplemental table 1:**
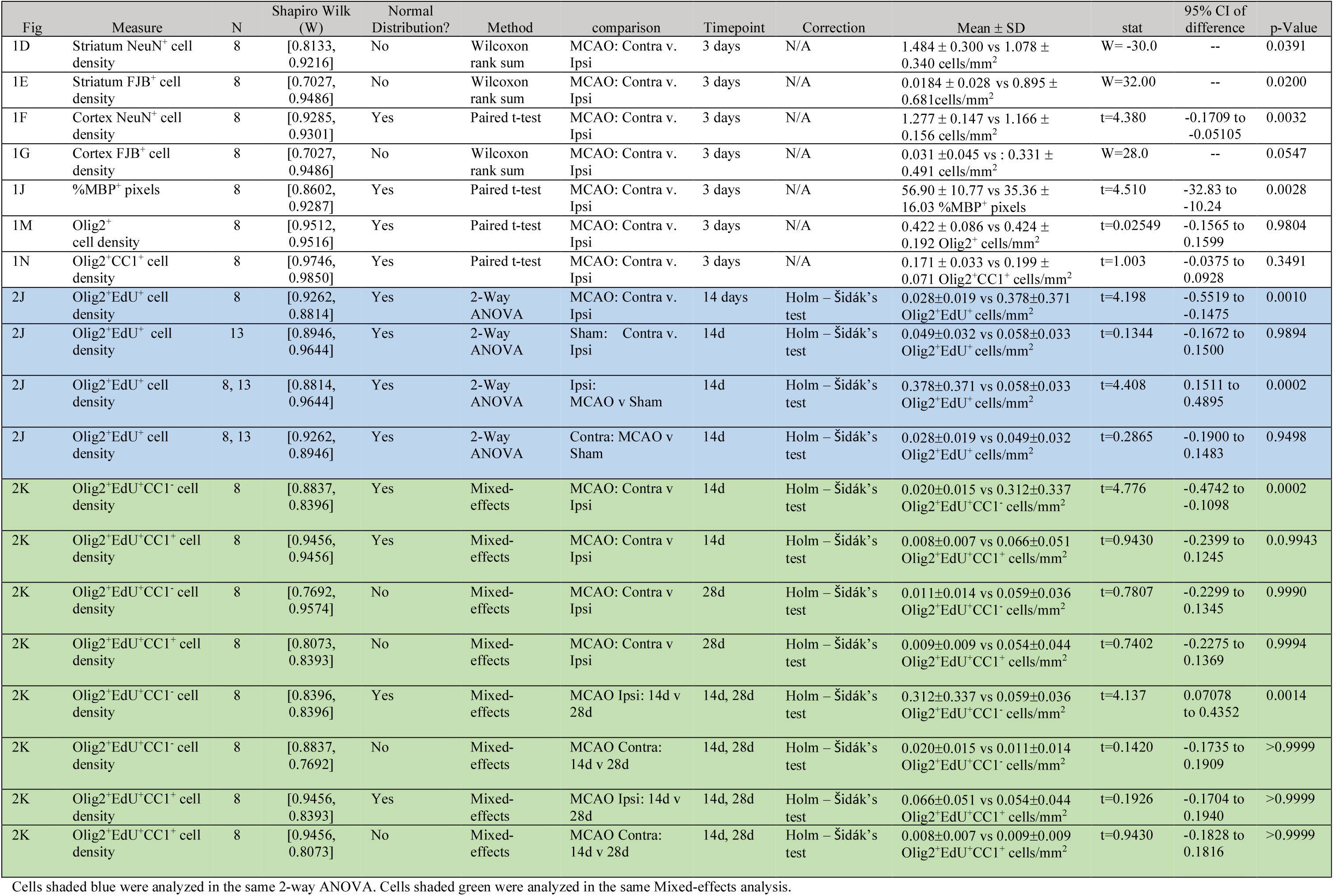
Statistical analysis for IHC data sets

**Supplemental Table 2:** General linear mixed model analysis of g-ratio, axon density and pathologic myelin.

| Fig | Measure | Comparison(N) | N value | Time point<br>(post-MCAO) | # mice<br>per group | mean±SE | estimate | SE of<br>estimate | df<sup>#</sup> | t-ratio | p-value* |
| --- | --- | --- | --- | --- | --- | --- | --- | --- | --- | --- | --- |
| 3E | Myelinated axon density | Contra(60) vs Ipsi(54) | Pencil fibers | 14d | 4 | 0.113±0.015 vs 0.116±0.015 myelinated axons/μm<sup>2</sup> | -0.00262 | 0.0156 | 165.2 | -0.168 | 0.9983 |
| 3E | Myelinated axon density | Contra(33) vs Ipsi(34) | Pencil fibers | 28d | 4 | 0.470±0.017 vs 0.399±0.017 myelinated axons/μm<sup>2</sup> | 0.07095 | 0.0198 | 165.6 | 3.581 | 0.0025 |
| 3E | Myelinated axon density | Contra: 14d(60) vs 28d(33) | Pencil fibers | 14d, 28d | 4 | 0.113±0.015 vs 0.470±0.017 myelinated axons/μm<sup>2</sup> | -0.35696 | 0.0227 | 12.7 | 15.704 | <0.0001 |
| 3E | Myelinated axon density | Ipsi: 14d(54) vs 28d(34) | Pencil fibers | 14d, 28d | 4 | 0.116±0.015 vs 0.399±0.017 myelinated axons/μm<sup>2</sup> | -0.28339 | 0.0228 | 12.8 | -12.422 | <0.0001 |
| 3F | g-ratio | Contra(838) vs Ipsi(719) | axons | 14d | 4 | 0.771±0.005 vs 0.772±0.005 | -0.00059 | 0.00308 | 4706.77 | -0.190 | 0.9976 |
| 3F | g-ratio | Contra(1693) vs Ipsi(1488) | axons | 28d | 4 | 0.798±0.005 vs 0.803±0.005 | -0.00485 | 0.00213 | 4713.50 | -2.277 | 0.1036 |
| 3F | g-ratio | Contra: 14d(838) vs 28d(1693) | axons | 14d, 28d | 4 | 0.771±0.005 vs 0.798±0.005 | -0.02772 | 0.00733 | 6.77 | -3.783 | 0.0288 |
| 3F | g-ratio | Ipsi: 14d(719) vs 28d(1488) | axons | 14d, 28d | 4 | 0.772±0.005 vs 0.803±0.005 | -0.03198 | 0.00737 | 6.96 | -4.337 | 0.0142 |
| 3G | Ratio abnormal axons | Contra (33) vs Ipsi (34) | Pencil fibers | 28d | 4 | 0.097±0.013 vs 0.129±0.015 | -0.0329 | 0.0137 | 58.8 | -2.406 | 0.0193 |

Results of the same shade fill (gray or white) were analyzed within the same glmm.

**Supplemental table 3:**
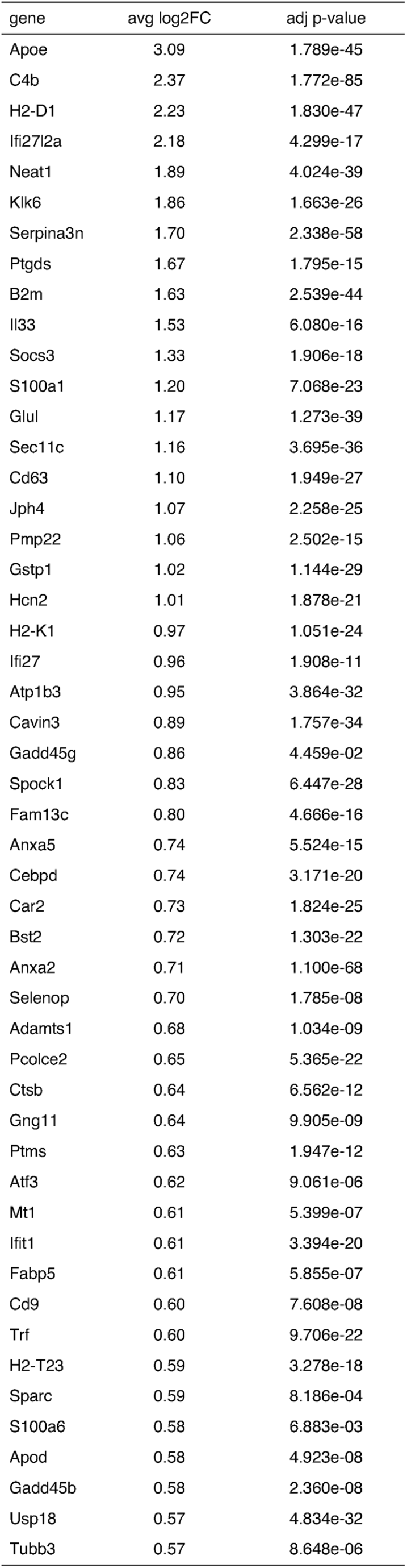
Top 50 genes upreglated in DOL cluster compared to Pre-OL2/MFOL/MOL

**Supplemental table 4:**
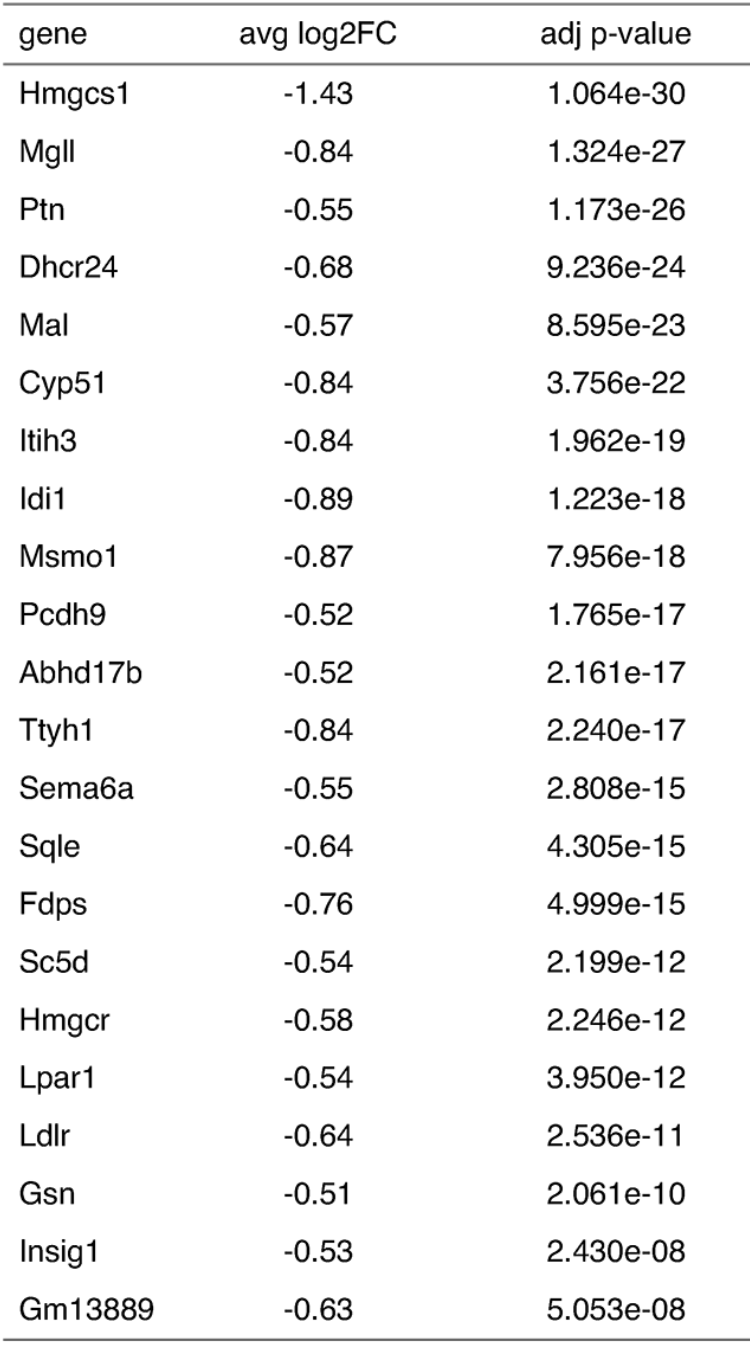
Top 50 genes downregulated in DOL cluster compared to Pre-OL2/MFOUMOL

**Supplemental table 5:** Marker genes of the DOL cluster

|  |  | fold change and adjusted p-value in Reactive cluster compared to each cluster |  |  |  |  |  |
| --- | --- | --- | --- | --- | --- | --- | --- |
|  |  | OPC | COP | Pre-OL1 | Pre-OL2 | MFOL | MOL |
| Inflammatory |  |  |  |  |  |  |  |
| Il33 | avg log 2FC<sup>†</sup> | 1.68 | 1.73 | 1.72 | 1.60 | 1.57 | 1.47 |
|  | p.adj | 1.44e-28 | 1.35e-21 | 3.76e-24 | 1.68e-01 | 2.82e-18 | 7.29e-07 |
| C4b | avg log 2FC<sup>†</sup> | 2.52 | 2.49 | 2.44 | 1.73 | 2.38 | 2.44 |
|  | p.adj | 3.14e-74 | 7.77e-45 | 1.07e-45 | 1.50e-07 | 2.15e-71 | 5.48e-56 |
| SerpinA3n | avg log 2FC<sup>†</sup> | 1.70 | 1.71 | 1.76 | 1.00 | 1.72 | 1.76 |
|  | p.adj | 5.44e-27 | 3.42e-21 | 1.01e-28 | 2.51e-02 | 2.06e-44 | 1.07e-37 |
| Klk6 | avg log 2FC<sup>†</sup> | 2.40 | 2.40 | 2.40 | 1.22 | 1.63 | 2.37 |
|  | p.adj | 1.68e-62 | 1.85e-37 | 2.74e-42 | 8.33e-01 | 3.85e-11 | 5.84e-55 |
| Other |  |  |  |  |  |  |  |
| Neat1 | avg log 2FC<sup>†</sup> | 3.19 | 3.14 | 2.95 | 1.81 | 1.74 | 2.14 |
|  | p.adj | 8.92e-79 | 5.54e-53 | 1.46e-52 | 6.41e-10 | 1.52e-32 | 3.87e-38 |
| Jph4 | avg log 2FC<sup>†</sup> | 1.37 | 1.53 | 1.60 | 1.57 | 1.24 | 0.83 |
|  | p.adj | 1.83e-37 | 1.92e-36 | 4.16e-48 | 2.49e-10 | 1.02e-33 | 2.48e-09 |

<sup>†</sup> avg log 2FC in Reactive cluster compared to each listed cluster

**Supplemental table 6:**
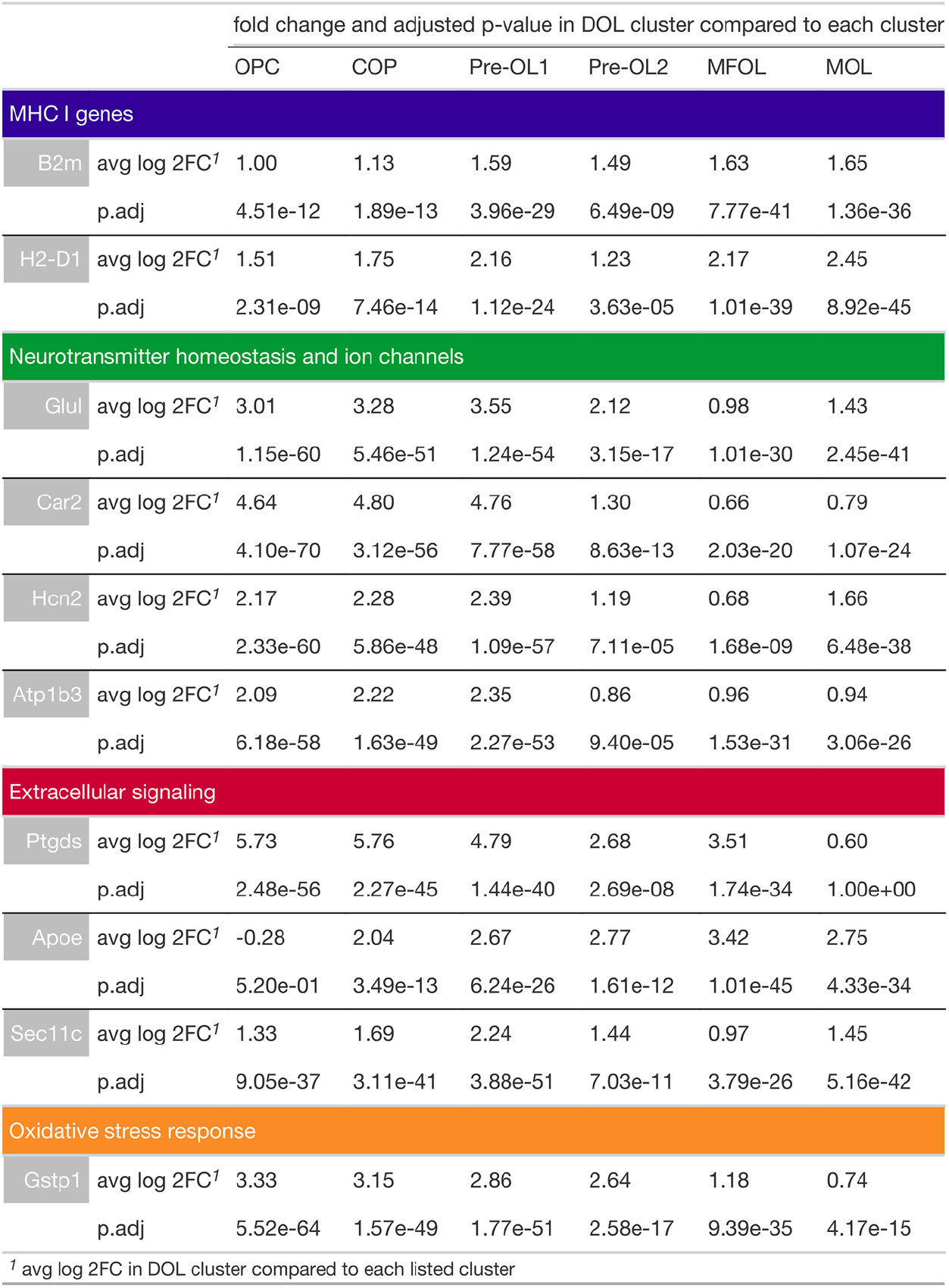
Genes differentially over-expressed in the DOL cluster

|  |  | fold change and adjusted p-value in DOL cluster compared to each cluster |  |  |  |  |  |
| --- | --- | --- | --- | --- | --- | --- | --- |
|  |  | OPC | COP | Pre-OL1 | Pre-OL2 | MFOL | MOL |
| MHC I genes |  |  |  |  |  |  |  |
| B2m | avg log 2FC<sup>†</sup> | 1.00 | 1.13 | 1.59 | 1.49 | 1.63 | 1.65 |
|  | p.adj | 4.51e-12 | 1.89e-13 | 3.96e-29 | 6.49e-09 | 7.77e-41 | 1.36e-36 |
| H2-D1 | avg log 2FC<sup>†</sup> | 1.51 | 1.75 | 2.16 | 1.23 | 2.17 | 2.45 |
|  | p.adj | 2.31e-09 | 7.46e-14 | 1.12e-24 | 3.63e-05 | 1.01e-39 | 8.92e-45 |
| Neurotransmitter homeostasis and ion channels |  |  |  |  |  |  |  |
| Glul | avg log 2FC<sup>†</sup> | 3.01 | 3.28 | 3.55 | 2.12 | 0.98 | 1.43 |
|  | p.adj | 1.15e-60 | 5.46e-51 | 1.24e-54 | 3.15e-17 | 1.01e-30 | 2.45e-41 |
| Car2 | avg log 2FC<sup>†</sup> | 4.64 | 4.80 | 4.76 | 1.30 | 0.66 | 0.79 |
|  | p.adj | 4.10e-70 | 3.12e-56 | 7.77e-58 | 8.63e-13 | 2.03e-20 | 1.07e-24 |
| Hcn2 | avg log 2FC<sup>†</sup> | 2.17 | 2.28 | 2.39 | 1.19 | 0.68 | 1.66 |
|  | p.adj | 2.33e-60 | 5.86e-48 | 1.09e-57 | 7.11e-05 | 1.68e-09 | 6.48e-38 |
| Atp1b3 | avg log 2FC<sup>†</sup> | 2.09 | 2.22 | 2.35 | 0.86 | 0.96 | 0.94 |
|  | p.adj | 6.18e-58 | 1.63e-49 | 2.27e-53 | 9.40e-05 | 1.53e-31 | 3.06e-26 |
| Extracellular signaling |  |  |  |  |  |  |  |
| Ptgds | avg log 2FC<sup>†</sup> | 5.73 | 5.76 | 4.79 | 2.68 | 3.51 | 0.60 |
|  | p.adj | 2.48e-56 | 2.27e-45 | 1.44e-40 | 2.69e-08 | 1.74e-34 | 1.00e+00 |
| ApoE | avg log 2FC<sup>†</sup> | -0.28 | 2.04 | 2.67 | 2.77 | 3.42 | 2.75 |
|  | p.adj | 5.20e-01 | 3.49e-13 | 6.24e-26 | 1.61e-12 | 1.01e-45 | 4.33e-34 |
| Sec11c | avg log 2FC<sup>†</sup> | 1.33 | 1.69 | 2.24 | 1.44 | 0.97 | 1.45 |
|  | p.adj | 9.05e-37 | 3.11e-41 | 3.88e-51 | 7.03e-11 | 3.79e-26 | 5.16e-42 |
| Oxidative stress response |  |  |  |  |  |  |  |
| Gstp1 | avg log 2FC<sup>†</sup> | 3.33 | 3.15 | 2.86 | 2.64 | 1.18 | 0.74 |
|  | p.adj | 5.52e-64 | 1.57e-49 | 1.77e-51 | 2.58e-17 | 9.39e-35 | 4.17e-15 |

<sup>†</sup> avg log 2FC in DOL cluster compared to each listed cluster

**Supplemental table 7:**
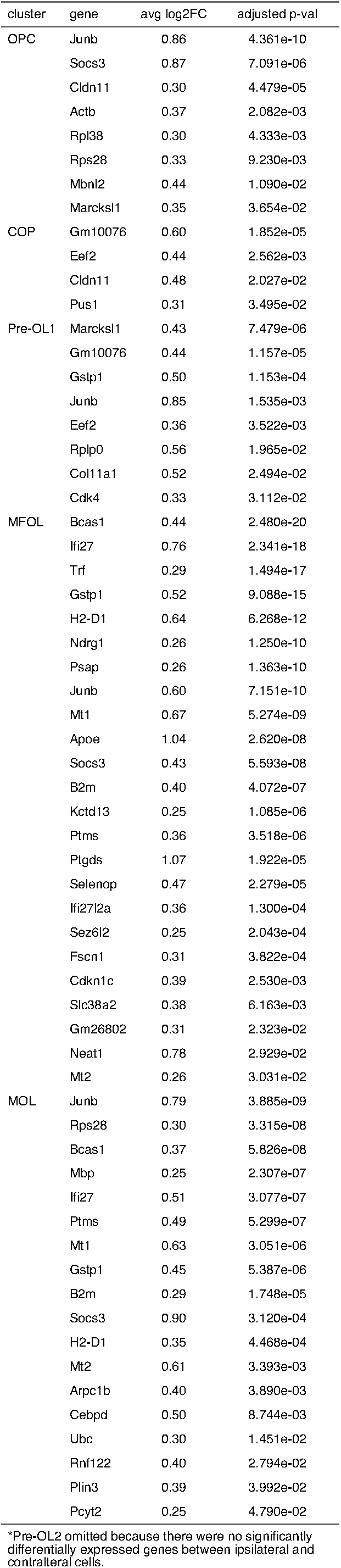
Genes significantly upregulated in ipsilateral oligodendrocytes*

**Supplemental table 8:**
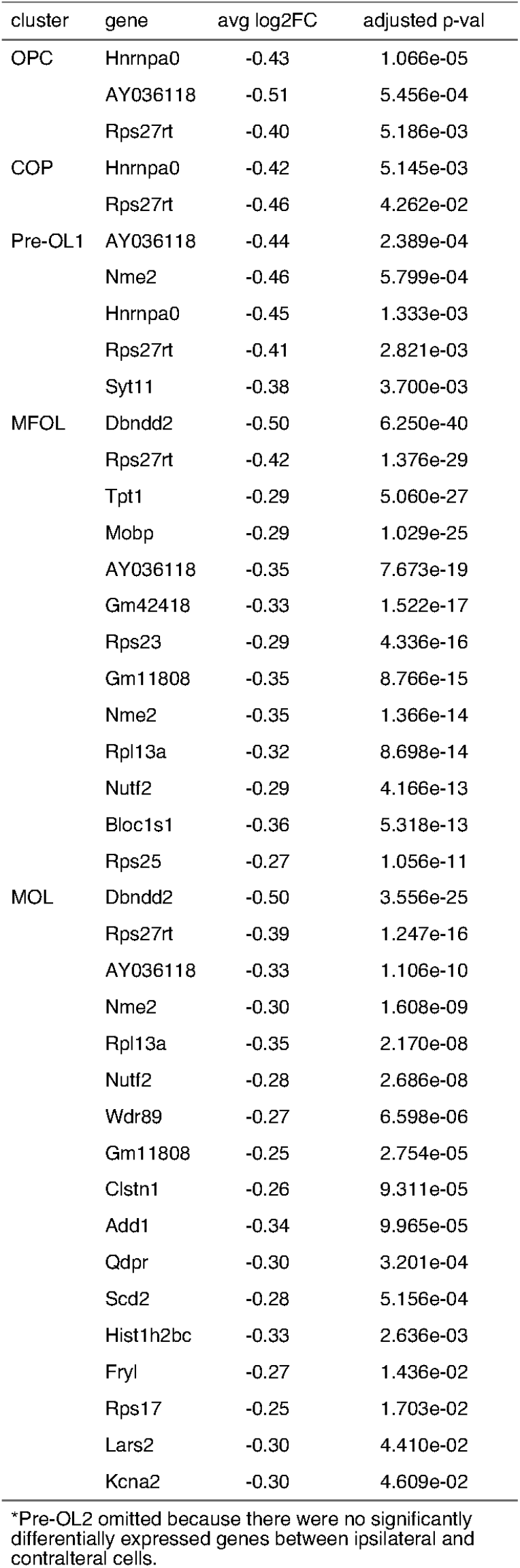
Genes significantly downregulated in ipsilateral oligodendrocytes *

